# Responses of grapevine cells to physiological doses of ethanol, among which induced resistance to heat stress

**DOI:** 10.1101/2024.08.31.610606

**Authors:** Alice Diot, Guillaume Madignier, Olivia Di Valentin, Anis Djari, Elie Maza, Yi Chen, Simon Blanchet, Christian Chervin

## Abstract

Grapevine naturally endures stresses like heat, drought, and hypoxia. A recent study showed very low oxygen levels inside grape berries, linked to ethanol content. Other studies have established the link between ethanol and tolerance to various stresses: heat stress, drought, and high salinity. The causes of such a tolerance are not well understood. In our study, three-week-old Gamay calli, *Vitis vinifera*, were characterised for their endogenous oxygen levels and endogenous ethanol concentration. Subsequently, a transcriptomic study of these cells was conducted, 6 and 24 hours after treatment with 1 mM ethanol. After 6 hours, ethanol addition led to 386 differentially expressed genes, with a notable upregulation of genes related to heat response, especially small Heat Shock Proteins (sHSPs). Further experiments showed that ethanol priming in grape cells or in *Arabidopsis* seedlings reduced pigment and electrolyte leakage under heat stress, respectively. This study supports the idea that ethanol priming helps protect plants against heat stress and provides a valuable RNA-seq dataset for further research into the underlying mechanisms, sHSPs playing a potentially crucial role in this adaptive response.

## Introduction

Hypoxia occurs in plant cells due to the insufficient availability of oxygen (Loreti and Perata, 2020). Two types of hypoxia can be distinguished: environmental hypoxia can be caused by specific environmental cues, while developmental hypoxia occurs in tissues and organs in well-oxygenated surroundings, under normal atmosphere oxygen availability (Jethva *et al*., 2022).

Environmental hypoxia is a condition periodically encountered by terrestrial plants, often in waterlogged soils (Daniel and Hartman, 2024). This phenomenon can also be induced by rising temperatures, as dissolved oxygen levels decrease with higher temperatures (Chapra *et al*., 2021) and increased microbial activity further depletes oxygen (Alkorta *et al*., 2017). Additionally, hypoxia occurs at high altitudes due to the reduction in oxygen partial pressure (Abbas *et al*., 2022).

Developmental hypoxia naturally occurs in bulky or dense tissues like seeds, fruits, teguments, and roots where oxygen diffusion is limited (Brazel and Graciet, 2023). It also occurs during rapid cell proliferation when the respiratory rate is intrinsically high: oxygen is consumed faster than it diffuses into the plant tissue (Geigenberger, 2003). For instance, Xiao *et al* (2018) reported very low levels of oxygen in grape berries during growth and late-ripening processes. Furthermore, developmental hypoxia has been observed in growing potato tubers (Geigenberger *et al*., 2000), in the embryos of faba bean (*Vicia faba*) and green pea (*Pisum sativum*) (Rolletschek *et al*., 2002), and in seeds during the germination process (Borisjuk and Rolletschek, 2009). Oxygen levels have been observed to decrease to hypoxic levels during rapid cell proliferation in specific organs, such as the shoot apical meristem and developing monocot anthers in *Zea mays* (Dukowic-Schulze and van der Linde, 2021), and during grapevine bud burst (Meitha *et al*., 2018).

Low oxygen concentrations induce cellular stress due to decreased ATP production, depletion of energy reserves, and accumulation of metabolic end-products notably ethanol (Jethva *et al*., 2022) through fermentative metabolism (Raymond *et al*., 1995). Ethanol also accumulates during particular plant developmental stages listed above: embryo and seedling development, fruit development, and maturation (Diot *et al*., 2024). It also accumulates in response to many other stresses including air pollution (sulphur dioxide and ozone exposure), water deficit, and freezing (Kimmerer and Kozlowski, 1982).

The impact of ethanol on plant physiology, particularly at low doses, is an emerging area of research as it could be a potential molecular marker of hypoxia (Diot *et al*., 2024). While high concentrations of ethanol are known to be toxic to plants, the few studies having investigated the effects of low physiological doses of ethanol on plants show beneficial outcomes, like resistance to heat stress (Matsui *et al*., 2022; Todaka *et al*., 2024), resistance to drought (Bashir *et al*., 2022) or resistance to high salinity (Das *et al*., 2022).

Matsui *et al*. (2022) have shown that exposure to physiological doses of ethanol modulates Heat Shock Protein (HSP) expressions in *Arabidopsis* seedlings. HSPs are molecules known to be expressed in response to a wide range of stresses, including thermal, chemical, and oxidative stress (Park and Seo, 2015). Among these HSPs, there is a subfamily called small heat shock proteins (sHSPs) which is proposed to act as molecular chaperones to protect other proteins from stress-induced damage (Waters and Vierling, 2020). The relationship between hypoxia and HSPs has already been studied, and it has been shown that the activation of HSPs is critical to adaptation to hypoxia and for enduring the oxidative stress of reoxygenation (Baird *et al.,* 2006). Chen *et al*., (2014) further showed that HSP70 accumulated more than 10 folds in the soybean plasma membrane in response to hypoxia stress. However, none of these studies established the link between ethanol, hypoxia and HSP. Moreover, most studies on ethanol investigate organisms’ response to “hammer” doses of this simple alcohol, i.e. non-physiological doses.

In the present study, we investigate the impact of ethanol at a physiological dose, 1 mM, on gene transcript expressions in grapevine cell cultures, as Xiao *et al*. (2018) have shown that hypoxic conditions occurred during grape berry development. Since the late 1970s, it has been widely accepted that plant cell cultures have numerous applications, among which serving as an excellent system for studying plant cell genetics, physiology, biochemistry, and pathology (Smetanska, 2008). However, there is a noticeable gap in the literature regarding the characterization of cell cultures grown on solid media. Indeed, most research focuses on plant cell suspension cultures (Hellwig *et al*., 2004; Sato, 2013), rather than on plant cell cultures grown on agar-solidified medium in Petri dishes (i.e., calli).

After characterising the physiological endogenous concentrations of oxygen and ethanol in these cell cultures, we carried out a whole-genome transcriptomic study using RNA-seq. Among the gene families up-regulated by a physiological dose of ethanol treatment, we identified genes encoding for HSPs. Following this observation, a series of metabolite leakage measurements confirmed the protective role of ethanol priming on grapevine cells exposed to heat shock, initially shown by Matsui *et al*. (2022) in lettuce leaves.

## Materials and methods

### Plant materials

Experiments were performed on a grapevine model. Cells from the berry of *Vitis vinifera* L. cv. Gamay Fréaux were sub-cultured in 55x15 mm Petri dishes using a previously published method (Triantaphylidès *et al*., 1993), with modifications concerning the culture medium. The composition of the growth medium used in this study is given in Supplementary Table S1. Parafilm was wrapped around the plates to prevent the medium and the calli from drying out. The plates were then placed in a climatic chamber (CRYO RIVOIRE, France) under constant conditions with a day/night photoperiod of 16h/ 8h, a temperature of 25°C/ 20°C and a light intensity of 150 µmol.m-1.s-1. Plates were moved every day within the chamber to ensure homogenous growth conditions.

A complementary experiment was performed on *Arabidopsis thaliana* seedlings var. Col-0. Seeds were sterilised using a 4.8% bleach solution for 10 minutes. Following sterilisation, the seeds were sown individually onto 35x15 mm Petri dishes containing MS/2 medium with 0, 0.1 or 1 mM ethanol (EtOH), using a sterile toothpick for placement. The Petri dishes were then sealed with micropore surgical tape (3M, 1.25 cm width, Germany) and stratified at 4°C for 24 hours. After stratification, the plates were transferred to a growth chamber (CRYO RIVOIRE, France) where the seeds were cultivated for 11 days under controlled conditions: light/dark, 25/20°C, 14h/10h and 150 µmol.m-1.s-1. Plates were moved every day within the chamber to ensure homogenous growth conditions.

### Characterization of cell cultures of *Vitis vinifera* L. cv. Gamay Fréaux

#### Callus development monitoring

Calli were grown for five weeks and photographed at 0, 7, 14, 21, 28 and 35 days in Petri dishes. The selected calli shown in the results were representative of the observed general development. For each time point, the average callus weight has been measured using a microscale. Calli grown for 14 and 35 days were harvested for cross-sections and photographed using the Axio Zoom.V16 microscope (ZEISS, Germany).

#### Endogenous ethanol measurements

For different stages, calli were frozen in liquid nitrogen. Each callus was ground to a fine powder and aliquoted at 200 mg/sample. The samples were then thawed in an Eppendorf 5415R centrifuge (Eppendorf, Germany) at 4°C and 13,600 g for 10 minutes. The supernatant was recovered and assayed for ethanol content using an enzymatic assay by measuring NADH absorbance at 340 nm (Ethanol kit, BioSentec, France)

#### Endogenous oxygen measurements

Dissolved oxygen in grapevine calli was measured using a Clark O_2_ microelectrode with a 25 μm diameter tip (OX-25; Unisense A/S, Denmark). The microelectrode was calibrated in 0% oxygen solution (0.1 M NaOH, 0.1 M C_6_H_7_NaO_6_) and in fully oxygenated water (273 μmol.L^-1^ at 22°C) as 100% O_2_ solution. Individual callus (equilibrated to room temperature) was placed on a motorised micromanipulator. The microelectrode was placed in the middle of the callus and [O_2_] profiles were taken in depth towards the centre of the callus at 300 μm increments. Oxygen readings were recorded using Unisense Suite software (Unisense A/S). Measurements were performed using 4 calli for each week of development.

### EtOH treatment and sampling for RNA sequencing

We selected the callus stage (i.e. 21 days, Fig. 1A) to have sufficient weight of plant material for the RNA extractions and because calli were in a healthy growing phase. Considering the endogenous ethanol levels in the callus at 21 days (406 ± 2 μM, Fig. 1C), we chose to treat our calli with 1 mM EtOH for the experiment. This treatment represents a 13% increase in ethanol concentration based on average callus weight and the volume of the treatment solution (see Supplementary Table S2). Due to the limited research on this subject, two sampling times were chosen: 6 hours and 24 hours post-treatment, to cover what we called short and long-term responses. Cell cultures were thus treated with either 100 µL of a control solution (fresh growth medium) or with 100 µL of a fresh growth medium containing 1 mM EtOH; and then placed back in their culture chamber for either 6 hours or 24 hours. Growth culture medium for this treatment purpose was prepared without agar. After the incubation time, each callus was then collected in 10 mL Falcon^TM^ tubes and immediately immersed in liquid nitrogen, then stored at -80°C.

**Fig. 1.**
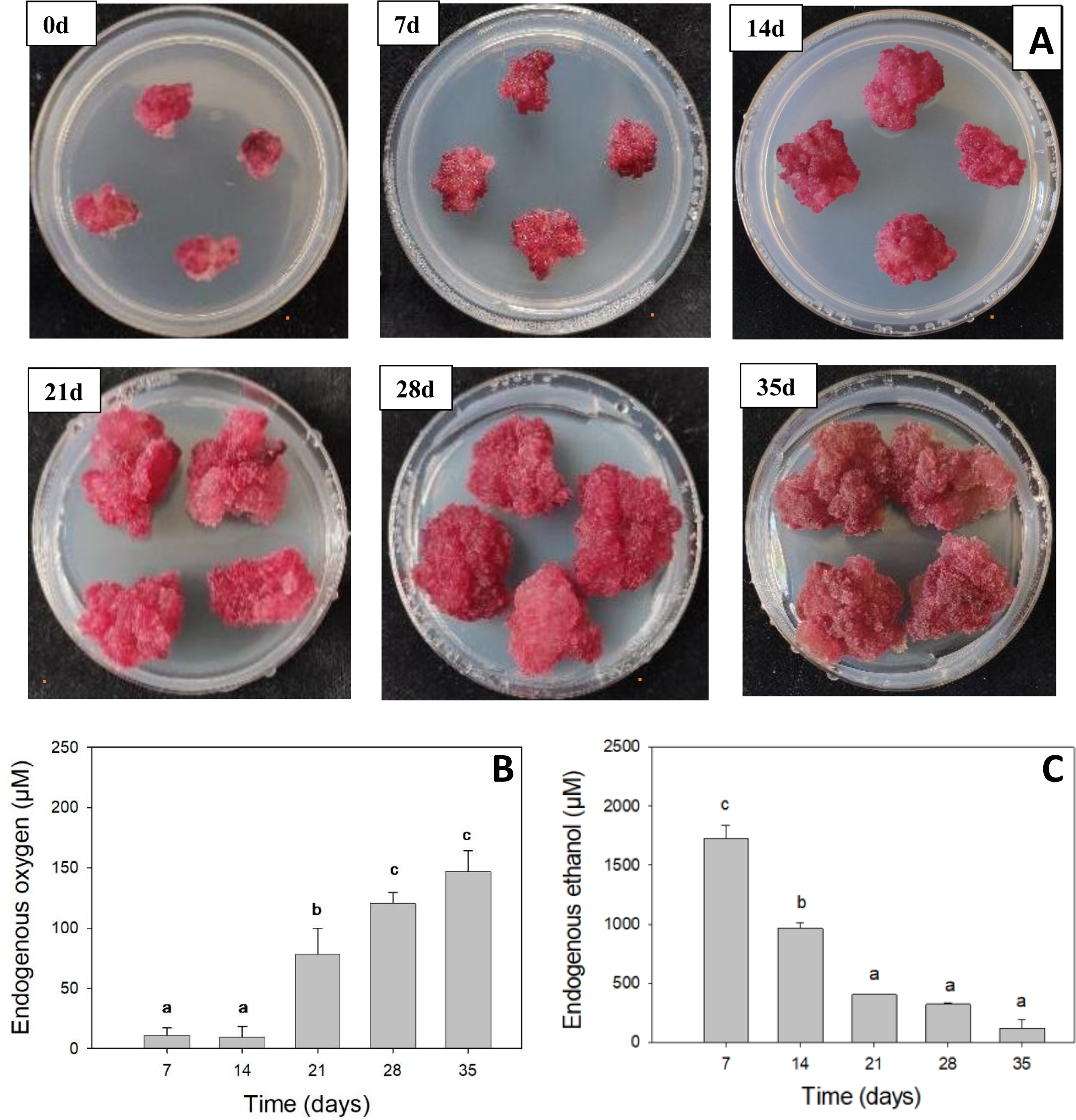
**A)** Photographs of Gamay calli at 0 day, 7 days, 14 days, 21 days, 28 days, and 35 days after subculturing onto Petri dishes. The calluses selected are representative of the general development observed. Scale: orange bar represents 1 cm. **B)** Dissolved O_2_ levels in callus at a distance of 2100 μm towards the callus centre as a function of development time (0, 7, 14, 21, 28 and 35 days), n = 4 biological replicates. **C)** Ethanol concentration as a function of development time (0, 7, 14, 21, 28 and 35 days), n = 3 biological replicates. Error bars represent standard error and different small letters indicate a significant difference at *p* < 0.05 by multiple comparisons using Fisher’s LSD for endogenous oxygen content and Tukey’s HSD for endogenous ethanol content.

### RNA extraction and purification

Frozen callus samples were homogenised to powder with the ball grinder Tissue Lyser II (Qiagen, Germany) by applying one cycle of 2 minutes at 300 Hz. Total RNA from individual calli was extracted using 160 mg of frozen powder with the Promega ReliaPrep™ RNA Tissue Miniprep System kit (Promega, France, Ref. Z6112) following the Promega - ReliaPrep™ RNA Tissue Miniprep System protocol, optimised for RNA extractions of grapevine cell cultures (supplementary protocol available on request from the corresponding author). RNAs underwent DNAse treatment with the Invitrogen™ ezDNase™ Enzyme kit to remove genomic DNA contamination from RNA preparations, following the ezDNase™ Enzyme kit protocol. For each sample, quantity and purity were first assessed by measuring the optical density (OD) at 260 nm and 280 nm with a NanoDrop® ND-1000 UV-Vis spectrophotometer (ThermoFisher, USA) and with a 1% (w/v) gel electrophoresis. Then, for each sample, RNA Integrity Number (RIN) was measured using an Agilent 2100 Bioanalyzer System microfluidic on-chip capillary electrophoresis device, following the RNA 6000 Nano kit protocol (Agilent Technologies, Germany). Samples with a RIN > 8.8 were selected and then aliquoted to obtain 20 μL RNA samples at 50 ng.μL^-1^.

### RNA-seq analyses

Selected RNA samples were sent to Novogene Bioinformatics Technology Co., Ltd (Beijing, China) for cDNA library construction and sequencing on the IlluminaNovaSeq 6000 platform (Illumina, San Diego, CA, USA). For each condition, three biological replicates were sent. Raw sequences corresponding to paired-end reads (2 x 150 bp) were treated as described by Althiab-Almasaud *et al*. (2021) and Chirinos *et al*. (2023) but adapted for the grapevine genome. The raw data discussed in this publication have been deposited in NCBI’s Gene Expression Omnibus (Edgar *et al*., 2002) and are accessible through GEO Series accession number GSE275842 (https://www.ncbi.nlm.nih.gov/geo/query/acc.cgi?acc=GSE275842). The reads were aligned on the *Vitis vinifera* L. cv. Chasselas annotated reference genome produced by Djari *et al*. (2024). The quality of these alignments is shown in Supplementary Tables S3A and S3B.

### Differential Expression and downstream analyses

First, we normalised the counts according to the transcript lengths to compare transcript expressions in downstream analyses. These normalised expressions can be understood as a relative estimation of transcript counts. The mean value of the biological replicates was calculated, giving as a result the estimated transcript counts for the given condition. Then, Differential Expression (DE) analysis was performed as described by Chirinos *et al*. (2023) with R software version 4.1.1 (https://www.r-project.org/) using the DESeq2 package version 1.30.1 with the default relative log expression (RLE) normalisation method, encompassing biases caused by library size and relative size of transcriptomes (Maza *et al*., 2013; Love *et al*., 2014). More precisely, DE analyses were performed as pairwise comparisons between all pairs of conditions (ex: “etha6h_1mM” versus “etha6h_0mM”). The false discovery rate (FDR) was controlled by the Benjamini–Hochberg method with an adjusted p-value (padj) < 0.05. Principal Component Analysis (PCA) was performed on estimated transcript counts with the plotPCA function of DESeq2 after a variance stabilising transformation.

### Identification and annotation of the differentially expressed genes (DEGs)

The functional annotation of the DEGs was retrieved from the VitExpress platform. When functional annotations were missing, gene ID codes were converted from “Vv” to the taxonomic ID of PNv4 “Vitvi” (Canaguier *et al*., 2017) using the GBFwebtool gene ID converter (https://www.grape.resources.gbfwebtools.fr/converter). The “Vitvi” codes were then used to obtain complementary annotations through MapMan Classification (https://mapman.gabipd.org/home) (Schwacke *et al*., 2019). If both annotations were missing, the “Vv” code was used to extract the corresponding gene sequence using the GBFwebtool “Extract Sequences” (https://www.grape.resources.gbfwebtools.fr/extractSequences) and was subsequently blasted in NCBI to find similar sequences and putative function (https://blast.ncbi.nlm.nih.gov/Blast.cgi).

### Gene Ontology Enrichment analysis

Gene ontology (GO) enrichment analysis was performed with R using the enrichGO function from clusterProfiler package version 4.10.1. Specifically, GO analysis was conducted separately on both the up-regulated and down-regulated DEG lists obtained from the comparison between EtOH-treated and control calli. Only DEGs with a |Log2FoldChange|>2 were selected.

### EtOH priming and heat-shock treatments

For *Vitis vinifera* cv. Gamay Fréaux cell cultures: 200 µL of liquid growth medium with EtOH at different concentrations (0, 0.1, 1, 10, 100 or 1000 mM) were poured onto three-week-old calli under sterile conditions. Petri dishes were then sealed with micropore surgical tape (3 M, 1.25 cm width, Germany) and returned to the culture chamber for 68 hours. After this incubation time, optimised by preliminary experiments, the heat shock (HS) was applied for 15 minutes in an incubator in the dark, with a peak temperature reaching 47°C (recordings of the heat stress peak can be found in Supplementary Fig. S1.). The Petri dishes were allowed to cool down for 20 minutes before returning to the growth chamber. Controls were treated with a control solution (liquid growth medium without EtOH) and were subjected to heat stress. Double negative controls were treated with a control solution (liquid growth medium without EtOH) and were not subjected to heat stress, but were kept in the dark for the duration of HS.

For *Arabidopsis thaliana* HS treatment, the protocol was adapted from Matsui *et al*. (2022) with the following modifications. *Arabidopsis* seedlings grown for 11 days with 0, 0.1 or 1 mM EtOH were washed with deionized water and placed in test tubes containing 1.8 mL of distilled water (15 shoots/tube). The heat stress was applied by placing the tubes in a water bath at 43 °C for 30 minutes and then cooling them down in a water bath set at 25 °C for 30 minutes. Control tubes did not undergo HS and were kept in a 25 °C water bath for 1 hour.

### Measurements of cellular leakage after HS treatment

To assess the cell damage caused by heat stress in *Vitis vinifera* and *Arabidopsis thaliana*, we selected cellular leakage as a key indicator of cell membrane integrity. Indeed, cellular leakage refers to the uncontrolled release of substances, such as ions, metabolites, or macromolecules, from the inside of a cell to its surrounding environment. This phenomenon typically occurs due to damage or disruption to the cell membrane (Jiang et *al.*, 2015), which normally serves as a selective barrier.

For *Vitis* cell cultures, cellular leakage measurements were assessed by measuring the coloured compounds released into the callus rinsing solution. The protocol was adapted from Ilík *et al*. (2018) with the following modifications: 72 hours after HS, calli were harvested, placed in 2 mL Eppendorf tubes, and weighed. To each tube, 1 mL of 100 mM mannitol rinsing solution was added to limit osmotic shocks. The tubes were then vortexed for 15 seconds and incubated at room temperature for 20 minutes on a rotator wheel, followed by centrifugation for 2 minutes at 16,100g. The absorbance of the supernatant was measured at wavelengths of 420 nm, 520 nm and 620 nm (Chervin *et al*., 2001; Lambri *et al*., 2015). The 100 mM mannitol solution was used as a blank.

We reported the results as relative absorbances, normalised to the HS controls (absorbance values set to 1) to account for variability between different batches.

For *Arabidopsis* seedlings, cellular leakage was assessed by the Index of Injury (Id) (Ilík *et al*., 2018), calculated from electrical conductivity measurements. The Id is a quantitative measure used to assess the extent of damage in plant tissues in response to stress conditions such as temperature extremes. It provides an indication of the degree of cellular or tissue damage, and evaluates the impact of various stressors on plant health. Based on a protocol adapted from Matsui *et al*. (2022), *Arabidopsis* seedlings grown for 11 days were washed with deionized water and placed in test tubes containing 1.8 mL of distilled water (15 shoots/tube). The heat stress was applied as described previously. Heat-stressed and control samples were slowly shaken overnight at 30 rpm, after which electrical conductivity was measured with a conductivity metre (Yangers, UK). The samples were then heated for 1h in a 100 °C water bath and shaken at room temperature for several hours, after which the electrical conductivity of the samples was again measured. The index of injury (Id) was calculated based on the protocol described for ion leakage analysis of *Arabidopsis seedlings* by Ilík *et al*. (2018).

### Anthocyanin content determination

This protocol was adapted from Vitrac *et al*. (2000), which investigated cell suspensions of *Vitis vinifera* (L.) and more notably those of cv. Gamay Fréaux Teinturier. Some modifications have been made: Three-week-old calli were treated with 100 µL of either a 1 mM EtOH or a control solution and were incubated for up to 16 days post-treatment in their growth chamber. Fresh callus tissues were collected, weighed, and placed into 1.5 mL Eppendorf tubes. Each tube received 500 µL of a solvent mixture composed of methanol and 0.32M HCl (85:15, v/v). The tissues were ground with a tube pestle for 1 minute, followed by incubation for 2 hours at 4 °C on a rotator wheel. After incubation, the tubes were centrifuged at 16,000g for 2 minutes. The supernatant was diluted to 1/50th of its original volume with the same solvent.

The absorbance of the anthocyanin extract was measured at 535 nm. Anthocyanin content was calculated using the molar extinction coefficient (log ε = 4.53) as reported by Vitrac *et al*. (2000). Methanol and 0.32M HCl (85:15, v/v) solvent was used as a blank.

### Total Polyphenol Index determination

The Total Polyphenol Index (TPI) is a measure used to quantify the total amount of polyphenolic compounds present in a sample. The TPI method relies on the absorption of benzene rings at 280 nm, a characteristic feature of polyphenols. This protocol was adapted from Cetó *et al*. (2012) with the following modification: on the same supernatant extracts as those prepared for anthocyanin determination, absorbance was measured at 280 nm using a quartz cuvette. The TPI for each sample was calculated as the absorbance multiplied by the appropriate dilution factor. Methanol and 0.32M HCl (85:15, v/v) solvent was used as a blank.

### Statistical analyses

All data in Figures 1 and 4 were analysed by ANOVA and multiple comparison tests, performed with Sigmaplot v.15 (Inpixon, USA). The confidence interval was set at 95% and the p-value is set at 0.05. Significantly different groups are indicated by small letters (a, b, c) on the graphs.

For RNA-seq data, the statistical analyses were performed using the R software, as specified in the previous paragraphs.

## Results

### Grapevine *Vitis vinifera* L. cv. Gamay Fréaux cell cultures are characterised by hypoxia and physiological ethanol levels in the µM to mM range

Figure 1A shows pictures of grapevine cell cultures as a function of development time over five weeks. The increase in callus size throughout their development is noticeable, but between days 28 and 35 of development, callus size varies only slightly (average callus weight over 5-week-development can be found in Supplementary Fig. S2). Furthermore, the typical colour of our study model, a bright pink to light red, is evident in the first four weeks of development. A noticeable colour change occurs between the 28th and 35th days of development: by the 35th day, the calli appear duller compared to those grown for 0, 7, 14, 21, and 28 days.

We then characterised the levels of oxygen in the grapevine calli during five weeks of development on growth media. Figure 1B shows dissolved O_2_ levels at a distance of 2100 μm towards the callus centre as a function of development time. We found that O₂ levels increase substantially and gradually over time of development, starting with concentrations of around 10 μmol.L^-1^ in 7-day-old calli to 150 μmol.L-1 in 35-day-old calli. Dissolved oxygen levels in calli grown for 7 and 14 days were statistically lower than in 35-day-old calli. Complete O_2_ profiles inside calli were measured and sections of 14-day-old and 35-day-old calli were observed, and results are shown and detailed in Supplementary Figure S3, and Supplementary Figure S4, respectively. Regarding the natural endogenous levels of ethanol in calli, we found a strong decrease in ethanol concentration as a function of development time, with an inverse relationship to oxygen levels (Fig. 1C). Ethanol concentration was significantly higher in 7-day-old calli than in 35-day-old calli, decreasing from 1726 ± 190 µM to 121 ± 99 µM, respectively.

Therefore, we can define the range from 100 µM to 1700 µM as the “physiological” ethanol concentrations present in cell cultures of *Vitis vinifera* L. cv. Gamay Fréaux under normal growth conditions.

### A small exogenous EtOH dose impacts the grapevine transcriptome

To investigate the impact of a physiological dose of EtOH on the *Vitis vinifera* transcriptome, we employed RNA sequencing (RNA-seq). Overviewing the transcriptional dynamics of the RNA-seq experiment, the Principal Component Analysis (Fig. 2A) revealed significant distinctions among the generated transcriptomes, with 74% of the variance associated with PC1 and 14% with PC2. The data consistently demonstrates a clustering pattern, wherein transcriptomes align closely according to their respective treatments, forming 4 distinctive clusters (0mM_6h, 1mM_6h, 0mM_24h, and 1mM_24h). This showed that biological variability was smaller than the differences between the four conditions and suggests a discernible impact of a low 1 mM dose of EtOH on the transcriptomes. There is more distance between the control and ethanol treatment at 6 hours than at 24 hours (Fig. 2A). Additionally, it is noteworthy that the 1mM EtOH treatment at 6 hours and at 24 hours show the greatest distance in the PCA plot, whereas the control solution at 6 hours and 24 hours generated the closest transcriptome groups. There were 5527 DEGs in the 6 hours comparison whereas there were 3755 DEGs in the 24 hours comparison (Supplementary Fig. S5).

**Fig. 2.**
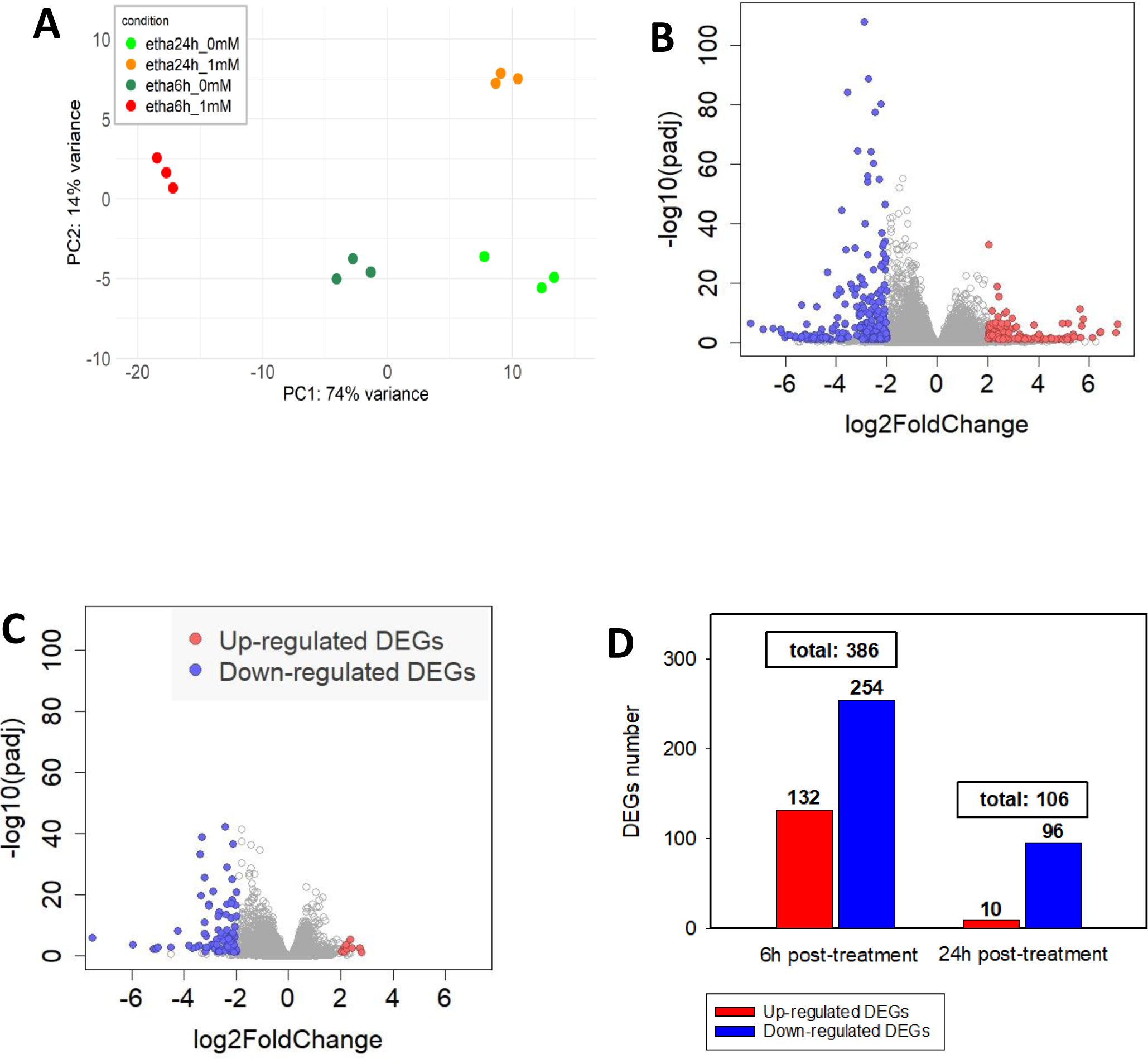
**A)** Principal Analysis Component of transcriptomic profiles of Gamay Fréaux grape cell cultures 6 and 24 hours after a control 0 mM (green and dark green) or 1 mM (orange and red) ethanol treatment. Every symbol represents one replicate. **B)** Volcano plot of 24655 genes for differential gene expression, at 6 hours post-treatment. **C)** Volcano plot of 24655 genes for differential gene expression, at 24 hours post-treatment. For both Volcano plots: cell cultures treated with 1mM EtOH versus control cell cultures. Scattered points represent genes: the x-axis is the Log2FoldChange for the comparison of EtOH treated cells vs control cells, whereas the y-axis is the negative log10 of adjusted p-values (where P is the probability that a gene has statistical significance in its differential expression). Differentially Expressed Genes (DEGs) that are significantly upregulated (Log2FoldChange > 2 and adjusted p-value < 0.05) are marked in red, while significantly downregulated genes (Log2FoldChange < -2 and adjusted p-value < 0.05) are marked in blue. Genes that do not meet both the significance threshold and one of the log2FoldChange cutoffs are shown in grey. **D)** Comparative Analysis of DEG number for the comparison 1mM EtOH versus control, at two time points. The DEGs shown in the figure are any genes with a p-adj < 0.05 and a Log2FoldChange > 2 for the up-regulated DEGs or Log2FoldChange < -2 for the down-regulated DEGs.

To further refine our analysis, we applied more stringent criteria to DEGs: we selected genes with a |Log2FoldChange|>2, in addition to requiring an adjusted p-value < 0.05 (raw results of the Differential Expression Analysis of the comparison 1 vs 0 mM EtOH at 6 hours are available in Supplementary Table S4). For the comparisons at both 6 and 24 hours, the DEG analysis revealed that there was a greater number of down-regulated DEGs (Fig. 2C, D). These results show a transcriptional trend towards inhibition, with most genes under-expressed in ethanol-treated calli. When visually comparing the number of DEGs between the two post-treatment durations, the volcano plot for the 6-hour time point shows a greater number of overall DEGs than the plot for the 24-hour comparison (386 vs. 106 genes respectively, see Fig. 2D).

These results indicate that even a light change in ethanol significantly impacts the grapevine transcriptome.

### EtOH treatment within the physiological range significantly upregulates small heat shock protein-related gene expressions

Considering the higher number of DEGs observed at 6 hours compared to 24 hours post-treatment, we decided to focus on the responses 6 hours after treatment. The complete lists of the 386 functionally annotated DEGs is available in Supplementary Table S5, sorted out in two sublists: Supplementary Table S5A provides the down-regulated DEGs (254 genes) and Supplementary Table S5B provides up-regulated DEGs (132 genes). Figure 3A shows the results of this GO enrichment for the up-regulated DEGs (p-adj < 0.05 and a Log2FoldChange > 2) while Figure 3B shows the GO enrichment for the down-regulated DEGs (p-adj < 0.05 and a Log2FoldChange < - 2) (the overall GO analysis results can be found in Supplementary Tables S6 and S7 respectively).

**Fig. 3.**
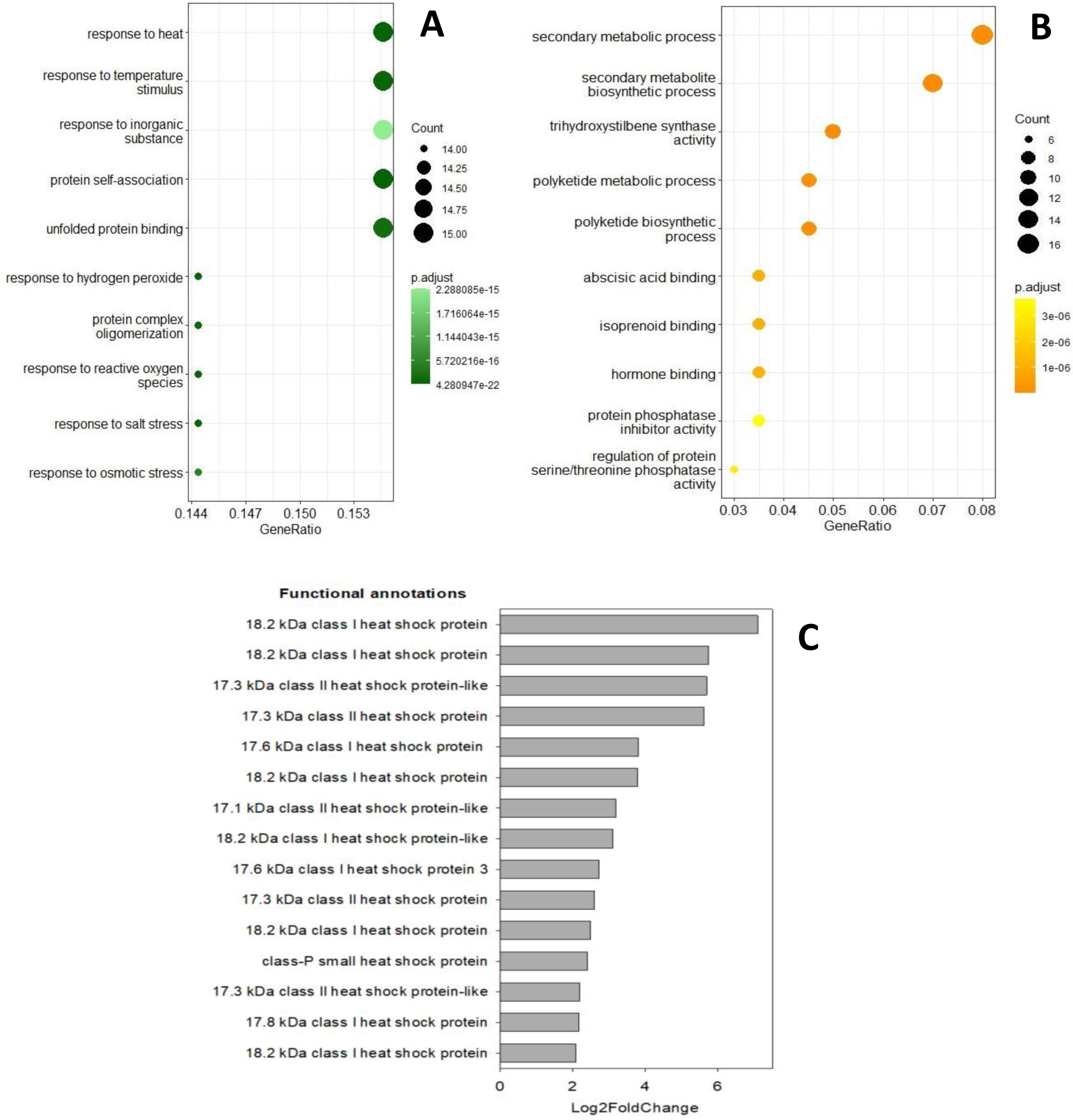
Dot plots of the Gene Ontology (GO) enrichment analysis of **A)** up-regulated DEGs and **B)** down-regulated DEGs for the comparison of 1mM EtOH versus 0mM at 6 hours post-treatment. DEGs used for the GO enrichment analysis are genes with p-adj < 0.05 and cutoffs as follows: Up-regulated DEGs-with a Log2FoldChange > 2 and down-regulated DEGs with a Log2FoldChange < -2. The size of the dots represents the number of genes in the significant DE gene list associated with the GO term and the colour of the dots represent the enrichment P-adjusted value. **C)** Details of the 15 up-regulated DEGs within the enriched GO class of “Response to heat “and ordered by their respective Log2 Fold Changes for a 1 mM ethanol treatment compared to control, at 6 hours post-treatment.

We found that genes annotated as “response to heat” (GO:0009408) and “response to temperature stimulus” (GO:0009266), “protein self-association” (GO:0043621) and “unfolded protein binding” (GO:0051082) were the most represented with a gene ratio of 0.158, among the 132 up-regulated DEGs (Figure 3A). For down-regulated genes, we found that genes annotated as “secondary metabolism process” (GO:0019748) and “secondary metabolite biosynthetic process” (GO:0044550) were the most represented among the 254 down-regulated DEGs, with gene ratios of 0.08 and 0.07 respectively (Fig. 3B).

Given these results, we had to examine the specific genes comprising the enriched families in both DEGs lists. First, we examined the specific genes comprising the up-regulated enriched family associated with the GO term “Response to heat”. This list includes 15 DEGs, presented in Figure 3C (their respective gene IDs, functional annotations, Log2FoldChange values, and adjusted p-values can be found in Supplementary Table S8). When looking at the functional annotations in Figure 3C, all the up-regulated DEGs are small Heat Shock Proteins (sHSP) and they are sorted out in two major groups: 9 genes are cytosolic class I sHSP and 5 genes are cytosolic class II sHSP. The last gene is a class-P sHSP. The Log2FoldChanges of these DEGs range from 2.09 to 7.09 and their associated p-adj values are very low (0.039 to 5.19^-12^), providing a high level of confidence that the ethanol treatment is responsible for this differential expression.

Then, we examined the specific genes comprising the down-regulated enriched family associated with the GO term “Secondary metabolism process”. This list includes 16 DEGs, presented in Supplementary Figure S6 (their respective gene IDs, functional annotations, Log2FoldChange values, and adjusted p-values can be found in Supplementary Table S9). When looking at the functional annotations in Supplementary Figure S6 we can notice genes involved in the general phenylpropanoid pathway like the pivotal phenylalanine ammonia-lyase (PAL), and genes involved in phenylpropanoid down-stream pathways. Indeed, genes involved in pathways for stilbenoid and flavonoid biosynthesis, such as stilbene synthase (STS) and chalcone synthase (CHS) are down-regulated. The Log2FoldChanges associated with these DEGs range from -2.02 to 4.90, being less extremes than the ones associated with the “Response to heat” genes.

### Ethanol-priming reduced heat-induced cell leakage

Following the observation of significant and highly up-regulated sHSPs revealed in the GO analysis, we looked for thermotolerance phenotypes in ethanol-primed calli after heat stress. The results are detailed in Figure 4. We observed that the double negative controls, which entails neither ethanol priming nor exposure to heat stress, exhibited the lowest absorbances at 420, 520 and 620 nm respectively (cyan bars, Fig. 4A-C), showing the basal cellular leakage in our experimental conditions. In contrast, cellular cultures exposed to both the control pre-treatment (0 mM EtOH) and HS demonstrated the highest absorbance values at all three wavelengths. Calli primed with EtOH solutions ranging from 0.1 to 10 mM, displayed a continuous decreasing trend in the absorbance values measured at 420 nm, 520 nm, and 620 nm, compared to the controls. The absorbances increased again within calli primed with 100 and 1000 mM EtOH solutions. These particular inverted bell-shaped curves are observed across all three monitored absorbances.

**Fig. 4.**
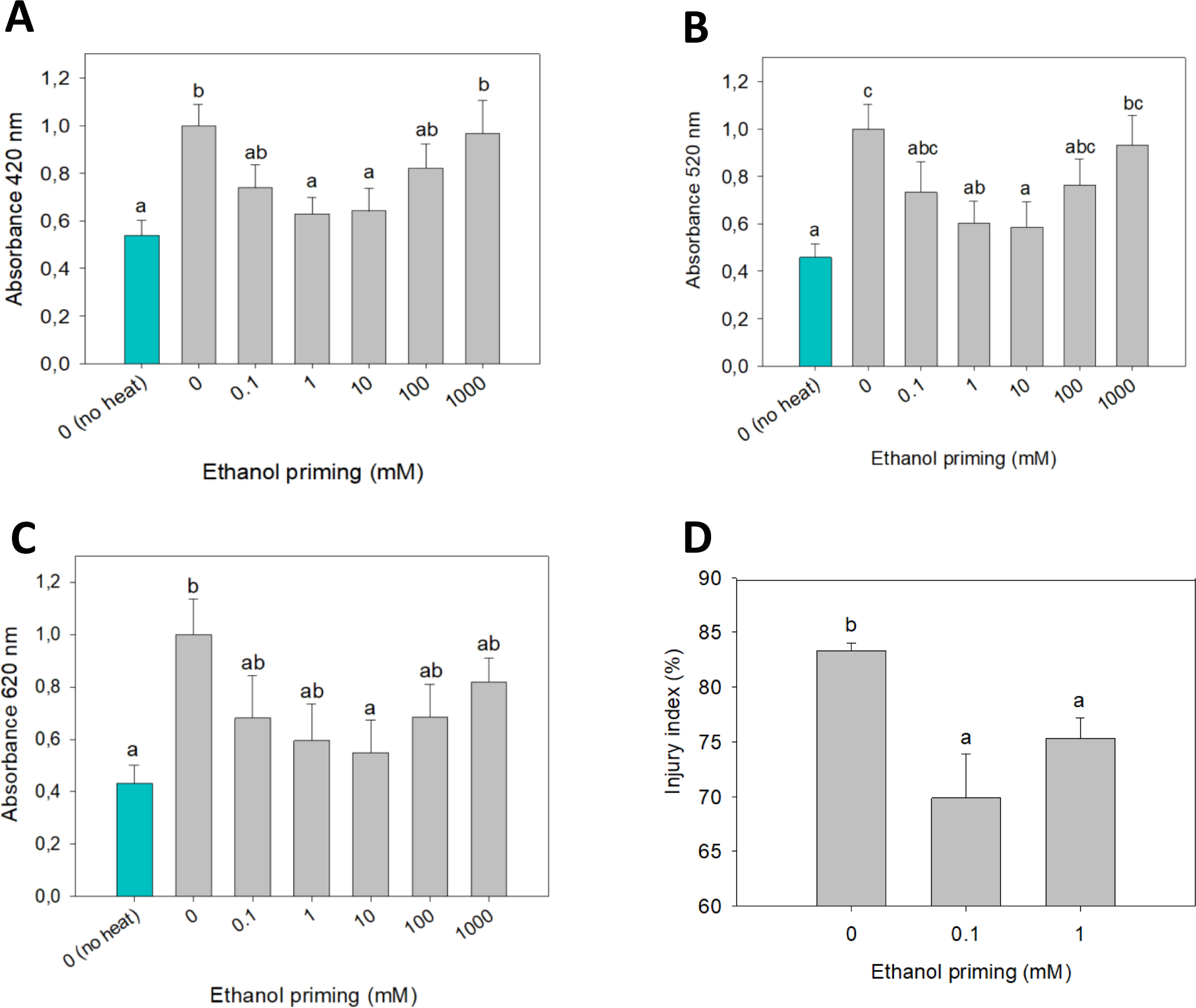
Damages induced by heat stress in *Vitis vinifera* Gamay Fréaux cell cultures and *Arabidopsis thaliana* Col-0 seedlings, as a function of pretreatments with various ethanol concentrations. Pigments released from Gamay cell cultures, absorbing at **A)** 420 nm, yellow range, **B)** 520 nm, red range and **C)** 620 nm, blue range, following a 47°C heat-stress (see material and methods for details). Results are means ± SE, n = 8 biological replicates, small letters indicate a significant difference (p < 0.05) between groups, multiple comparisons by Fisher’s LSD. **D)** Injury index of Arabidopsis seedlings based on ion leakage measurements following a 43°C heat-stress (see material and methods for details). Results means + SE, n = 7 biological replicates, small letters indicate a significant difference (p < 0.05) between groups, multiple comparisons by Tukey’s HSD.

Overall, calli primed with concentrations of 1 and 10 mM showed strong reductions in absorbance around 35-45% compared with the HS-control. These results show that priming grapevine cellular cultures with low physiological doses of ethanol (i.e. 0.1 to 10 mM) reduced the cellular compound leakage after exposure to a mild heat shock. Moreover, a complementary experiment was conducted on *Arabidopsis thaliana* seedlings to confirm that low doses of EtOH could limit cellular leakage in another plant’s tissues and in a different plant species. Figure 4D presents ion leakage for *Arabidopsis thaliana* seedlings. We found that in control seedlings, the injury index at 83% was significantly higher than in seedlings treated with 0.1 and 1 mM EtOH, which were at 69% and 75% respectively. This corresponds to a significant reduction in the injury index by 14% for the 0.1 mM ethanol treatment and 8% for the 1 mM ethanol treatment compared to control. This result shows that priming *Arabidopsis* seedlings with low physiological doses of ethanol (i.e. 0.1 and 1 mM) significantly reduced the ions released after exposure to a mild heat shock, and further reinforces the results described for the Gamay cell cultures.

### Ethanol-priming does not impact polyphenol content in grapevine calli

As shown in Figure 3B, the GO terms ‘Secondary metabolic process’ and ‘secondary metabolite biosynthetic process’ are significantly enriched among the 254 down-regulated DEGs, many of which are linked to polyphenol synthesis (see Supplementary Fig. S6). Given this significant down-regulation observed in secondary metabolism, and particularly the down-regulated PAL and CHS transcripts, we investigated polyphenol content in ethanol-primed calli. To assess this further, we monitored anthocyanin and total polyphenol content as a function of time in grapevine cell cultures, following an initial 1 mM EtOH treatment. The supplementary Figure S7 illustrates the changes in anthocyanin concentration and total polyphenol index in cell extracts. Both graphs show an increasing trend as a function of time, however, there was no significant difference in anthocyanin concentrations (Supplementary Fig. S7A) or in the total polyphenol index (Supplementary Fig. S7B) between the ethanol-primed and control treatments.

## Discussion

### Grapevine cell cultures, a model experiencing hypoxia and ethanol accumulation

A recent study by Xiao *et al (*2018) showed low levels of oxygen in grape berries and even anoxia in the centre of the berries. Our grapevine cell culture model shows the same trend, with developmental hypoxic to anoxic oxygen levels and inversely correlated ethanol concentrations in the calli, especially when they undergo rapid growth in the first 2 weeks after subculturing (Fig. 1B,C). This rapid growth is associated with a high rate of cell division and high cellular oxygen consumption in most plant tissues (Raymond *et al*., 1995). The increase in oxygen levels in calli, as they age, is probably linked to a decrease in their growth and metabolic rates. This slower growth allows oxygen to diffuse faster than it is consumed by the cells. Additionally, after five weeks of growth, cross-sections (Supplementary Fig. S4) revealed a heterogeneous structure with invaginations and gaseous interstices within the callus, further facilitating air and oxygen diffusion. A close relationship exists between oxygen and ethanol levels in plants, in which decreased oxygen levels result in increased ethanol concentrations, due to the shift from aerobic to anaerobic respiration (Bui *et al*., 2019). Thus, as dissolved oxygen levels increase over callus developmental time, it is logical to observe a corresponding decrease in ethanol levels. Therefore, we can conclude that our cell culture model naturally experiences developmental hypoxia, leading to ethanol accumulation. This validates the cell culture model for studying the impact of small changes in ethanol concentrations. These latter could be related to climate change, as rising temperatures in this context further reduce dissolved oxygen levels in water and organisms, while increasing respiration rates. These changes lead to hypoxic metabolism in certain organisms and tissues, and to ethanol production. The question arises whether ethanol serves as a signal prompting organisms to adapt, or if it is simply an end-product of hypoxic metabolism (Diot *et al*., 2024).

### Small exogenous application of EtOH triggers a fast response in the grapevine transcriptome and cells seem to perceive it as a stress

We present evidence that small changes in ethanol concentration generate quick responses at the transcriptomic level in Gamay cell cultures. On the PCA plot (Fig. 2A), the closely grouped transcriptomes based on their respective treatments and the high explained variance demonstrate the significant impact of ethanol on gene transcripts. The sum of the variances (PC1 + PC2), which accounts for 88% of the total variance, indicates that the two-dimensional plot effectively represents the impact of the ethanol treatment and the two sampling time points (6 and 24 hours). It is possible that the highly controlled *in vitro* culture environment and the use of identical cloned cells limit the biological variability in our experimental design. The fact that Gamay cell cultures can sense very small changes in ethanol concentration (1 mM) is evidenced by the 386 DEGs identified at 6 hours (Fig. 2D), suggesting that a small amount of ethanol is quickly detected by the grapevine cells and prompts a strong and immediate transcriptional response. The reduction in the number of DEGs to 106 at 24 hours post-treatment (Fig. 2D) shows that the transcriptional response diminished over time. This result is further reinforced by the decreased number of DEGs with a high Log2FoldChange after 24 hours compared to 6 hours (Fig. 2C versus Fig. 2B). This trend suggests that the transcriptional response to low doses of ethanol is not only strong and immediate but also relatively transient. The general trend of downregulation of the transcriptomic dynamics when comparing 1 mM EtOH to control treatments (Fig. 2D), at 6 and 24 hours post-treatment, is often observed in plant responses to stresses. Indeed, Ul Haq *et al*. (2019) showed that a plant’s common strategy to cope with stressful situations is reducing the synthesis of normal proteins, while enhancing the production of stress-related proteins, such as HSPs. This trade-off is clearly illustrated at the transcriptomic level in our study: Gamay calli downregulate genes involved in secondary metabolism (Fig. 3B) while simultaneously overexpressing a set of genes related to stress proteins following the addition of exogenous ethanol (Fig. 3A). This GO analysis revealed that cell sHSP genes were upregulated in calli treated with 1 mM EtOH compared to the control at 6 hours (Fig. 3A). sHSPs function as early responders to assist cells in managing proteins destabilized by stress conditions (Janowska *et al*., 2019), and their induction represents a rapid and intense emergency response (Parsell and Lindquist, 1993). This finding aligns with our RNA-seq data, as we observed that, just 6 hours after treatment with 1 mM EtOH, some sHSP genes were expressed more than 64 times than in the control (Fig. 3C). Our study suggests that a minor change in ethanol is sensed by the cells as a stress, prompting them to initiate adaptive responses.

### Responses of grapevine cells to physiological doses of ethanol: upregulation of small heat shock protein transcripts

Our results show that the genes associated with the GO terms “Response to heat” and “Response to temperature stimulus” are the most statistically overrepresented 6 hours after a 1mM EtOH treatment, notably the sHSPs (Fig. 3C).

Firstly, the ability of ethanol at physiological doses to stimulate sHSP synthesis has already been observed in the animal kingdom. Indeed, some studies showed that ethanol at 50 mM induced sHSP genes in mouse embryonic neural stem cells (Choi *et al*., 2011) and EtOH at 60 mM increased the levels of HSP27 in mouse cortical neurons (Pignataro *et al*., 2007).

<u>sHSPs:</u> HSPs are classified into five major subfamilies based on their molecular weights: Hsp100, Hsp90, Hsp70, Hsp60, and small heat shock proteins (sHSPs) (Sun and MacRae, 2005). These latter are differentiated from the other HSPs by their smaller molecular weight -typically, plant sHSPs range between 15 and 30 kDa-, and are the most abundant in plants (Vierling, 1991). The sHSPs are expressed during the normal developmental process of plants (Lindquist and Craig, 1988), and are known for their response to a wide range of environmental stresses, including heat, cold, drought, salinity, and oxidative stress; and it seems that the high diversification of plant sHSPs reflects a molecular adaptation to stress conditions. As it is clear that sHSPs have evolved to be sensitive to small changes in pH or in metal ions near physiological concentrations (Janowska *et al*., 2019), it would be logical that they are also sensitive to other small changes near-physiological state like, in our case, small changes in ethanol concentration. The small HSPs are particularly renowned for their crucial role in maintaining proteostasis when expressed in response to cellular stress in plant systems (Haslbeck and Vierling, 2015). Indeed, they specialized in preventing protein aggregation by binding to partially unfolded proteins under stress conditions (McLoughlin *et al*, 2016; Janowska *et al*., 2019; Lee *et al*., 1995, 1997), differentiating from other larger HSPs in their role, as sHSPs do not typically refold proteins. Besides this activity, sHSPs can function as membrane stabilisers and ROS scavengers or act synergistically with antioxidant systems. Overall, they play a key role in maintaining membrane fluidity and permeability under some stress (Aghdam *et al*., 2013). However, ethanol was shown to modify the fluidity of phospholipid bilayers (Patra *et al*., 2006), possibly creating a link between ethanol, membrane organization, and the induction of sHSPs.

Class CI and CII sHSPs:

In Figure 3C, the key genes attracting our attention were sHSP genes encoding for class CI and CII subfamilies. These are the two most abundant classes of plant cytosolic-localised sHSPs. CI sHSPs typically comprise the largest family of sHSP genes in higher plants (Waters, 2013) while the CII gene family is typically smaller. Basha *et al*. (2010) examined the structural and functional differences between the CI and CII subfamilies. They reported that both the CI and CII subfamilies have chaperone activities and they were effective at preventing irreversible aggregation, illustrating their role in the prevention of future damages made to the cell. Therefore, it is not so surprising that 6 hours after the EtOH treatment, we observed a significant up-regulation of these protein transcripts. Thus, we can hypothesize that the plant may use very small changes in ethanol to prevent further damage due to stresses. Indeed, ethanol has been shown to induce sHSP in soybean (Kuo *et al*., 2000) and in rice (Guan *et al*., 2004), even if the ethanol doses tested were 5% (v/v) corresponding to a concentration of nearly 1M, which can be considered like a “hammer” dose.

### Small changes in ethanol concentration confer thermotolerance to grapevine calli by reducing cellular leakage, a trend confirmed by experiments with *Arabidopsis* seedlings

Given that our plant model is a culture of cells rich in anthocyanins (Guan *et al*., 2016; Kong *et al*., 2021), we drew inspiration from studies demonstrating the release of pigments following heat stress (Lambri *et al*., 2015). Zhang *et al*. (2005) showed that heat treatment (38 °C for 10 h) induced injuries in the ultrastructure of *Vitis* mesophyll cells, then linked to increase in conductivity associated with membrane leakage, and also linked to malondialdehyde increase. In oenology, the postharvest heat treatment of grape berries has been known for a long time to increase berry skin cell damage and facilitate the release of anthocyanins (Lambri *et al*., 2015). These latter authors showed that heating grape berries at various temperatures from 60 to 80°C for various times 3 to 60 min, increased the absorbances at the main three wavelengths defining red wine colour intensity: 420, 520 and 620 nm, the main red pigments being anthocyanins (Lambri *et al*., 2015). In our study, after preliminary experiments, we chose a rather mild heat treatment (Supplementary Fig. S1) in order to illustrate differences induced by ethanol priming. In our Figure 4A, 4B and 4C, significant differences in pigment leakage indicate that ethanol priming protected grapevine calli from damage due to exposure to heat stress, compared to control (“0”). This protective effect was also shown in a different plant species. Indeed, significant differences in electrical conductivity indicate that ethanol priming also protected *Arabidopsis* seedlings from damage due to exposure to heat stress (Fig. 4D), compared to control (“0”). In other words, ethanol priming effectively reduced cell leakage caused by heat treatment, demonstrating that a low dose of ethanol enhances tolerance to heat stress, and suggesting that the ethanol-induced-sHSPs are related to the acquisition of thermotolerance. Increasing data suggest a strong correlation between sHSP accumulation and plant tolerance to stress (Liu *et al*., 2012; Wang *et al*., 2004), like our data. These latter authors suggest that the protective effect involves membrane stabilization. According to Ilík *et al*. (2018), heat acclimation is connected to an increase in the degree of saturation of fatty acids of membrane lipids, which in turn leads to increased rigidity of cell membranes and increase in their thermostability. A search for desaturase-related-genes in our RNA-seq data did not bring evidence that ethanol priming is regulating this type of enzyme, at 6 or 24 hours post-treatment. Nevertheless, Tsvetkova *et al*. (2002) showed that association between sHSPs and membranes may constitute a general mechanism that preserves membrane integrity during thermal fluctuations.

Plants have evolved many mechanisms for adaptation to dynamic and adverse environmental conditions. In addition, they have acquired a form of ‘stress memory’ in that they retain their response to an initial stress, which primes their response to a second stress, allowing them to respond more rapidly and strongly (Savvides *et al*., 2016). One adaptation is their ability to produce a variety of compounds that induce the expression of stress-related genes, whose encoded products protect cellular components from the adverse impact of abiotic stress (Sako *et al*., 2020). This phenomenon, called chemical priming, is evident in our results, as cell cultures that underwent ethanol priming showed better protection against mild heat stress. Todaka *et al*. (2024) demonstrated that a 20 mM ethanol treatment enhanced heat stress tolerance in tomatoes, and highlighted the potential of ethanol-based chemical priming as a technology to mitigate heat stress damage in crops. Crosstalk between heat and ethanol stresses, and heat and ethanol tolerance, has already been shown in several species and seems to be a conserved trait across kingdoms. In plants, this has been shown by Matsui *et al*. (2022). And in other kingdoms, HSP induction correlates with induced tolerance to high concentrations of ethanol: in mammals, Chinese hamster (Li, 1983); in microorganisms, *Saccharomyces cerevisiae* (Plesset *et a*l., 1982).

One point left to discuss is the timeframe of the responses. The sHSPs expression is often part of an immediate stress response (Hu *et al*., 2022) but their synthesis is not maintained through time (Altschuler and Mascarenhas, 1982). Our results reflect this, as no sHSP genes were up-regulated 24 hours post-ethanol treatment when compared to control. However, 68 hours after ethanol priming, thermotolerance phenotypes were observed (Fig. 4A-C). Vierling (1991) reported that pea plants exposed to heat stress accumulated chloroplastic sHSPs and class CI HSPs, even after the stress had ended. The half-life of these proteins was found to be around 52 hours for chloroplastic sHSPs and 37 hours for cytoplasmic HSPs. This shows that although the expression of small heat shock proteins is transient, the sHSPs themselves are quite stable. The persistence of these HSPs suggests that their participation in recovery processes could be as important as their possible role during the stress period itself (Vierling, 1991). However, the authors of the present study have to report that priming is effective in mitigating heat stress when the stress is mild. Indeed, when the stress exceeds a certain threshold (higher than sublethal heat shock), the priming effect diminishes, and the treated cells and *Arabidopsis* seedlings exhibit worse leakage compared to the control (data not shown).

### Responses of grapevine cells to physiological doses of ethanol: downregulation of key polyphenol-related transcripts, without effects on anthocyanin and polyphenol contents

As previously discussed, the general trend in the downregulation of secondary metabolism is a classical plant response to stresses (Ul Haq *et al*., 2019; Bilgin *et al*., 2010; Liu *et al*., 2012). However, no significant difference was observed between EtOH treated and untreated calli regarding the content of anthocyanins and the content of total polyphenols (Supplementary Fig. S7). We have two main hypotheses to explain the difference between the transcriptomic and the physiological levels. First, it could be that because the anthocyanin and polyphenol contents are so important in Gamay Fréaux calli, that the transient downregulation of the enzymes involved in the biosynthesis of polyphenols results in a too small physiological change. Indeed, the Gamay Fréaux is a teinturier grape and stands out among grape cultivars due to its rare and unique red-fleshed berries as well as red skin, which is uncommon compared to the majority of grape cultivars having red skin colour but white-coloured flesh (Guan *et al*., 2016). Second, the polyphenol pathway is complex with numerous proteins involved. The fact that the expression of some of these protein transcripts is inhibited does not ensure that polyphenolic compounds accumulate at different levels.

### Plant response to ethanol displays hormesis characteristics and ethanol could be a potential messenger of hypoxia and other stressful conditions

Because we saw the sHSPs subfamily being extremely overexpressed in the RNA-seq experiment and in order to examine a physiological response of grapevine cell cultures to low doses of ethanol, we assessed the cell damages resulting from a heat shock in calli primed with ethanol at different concentrations. The results (Fig. 4A-C) show a clear biphasic dose-response curve for the absorbances tested, typical of an hormesis effect. Hormesis is a phenomenon observed in biology where exposure to low doses of a stressor or toxin results in a beneficial effect on an organism, contrary to the expected harmful effect at higher doses (Calabrese and Baldwin, 2002; Calabrese and Baldwin, 2003). This concept challenges the traditional linear dose-response model by proposing U-shaped or inverse U-shaped response curves (Erofeeva, 2022). According to Calabrese and Baldwin (2002; 2003), hormesis is considered an adaptive response of organisms to stimuli (environment, chemicals, …). At low doses, the stressor triggers adaptive responses within cells and organisms, such as increased antioxidant production (Wang *et al*., 2023) and repair mechanisms (Scott, 2008), which enhance resilience and improve overall health. Calabrese and Baldwin (2002; 2003) suggest that this stimulatory response could be the result of compensatory biological processes, following an initial disruption in homeostasis, i. e. in our study, the slight change in ethanol concentration. Although there is no single biological mechanism that can explain hormesis across plants, Agathokleous *et al*. (2020) suggest that certain general principles regarding the mechanisms of hormesis across different species and stress-inducing agents can be identified. These authors note that low doses of stress induce a mild increase in the production of reactive chemical species, hormones, and enzymatic and nonenzymatic antioxidants, as well as upregulation of proteins (e.g., heat shock proteins); and that these responses lead to reduced toxicity if plants are subsequently exposed to severe stress, within a preconditioning framework. Our study shows both an increase in heat shock proteins transcripts (Fig. 3B) and reduced HS toxicity (Fig. 4) after an ethanol treatment. This hormesis effect of ethanol has already been seen in a study by Middleton *et al* (1978) on its impact on the architecture of pea roots, where the mean root length was increased by an ethanol concentration as low as 0.002 ml.L^-1^ (equivalent to 0.0002% or 0.034 mM). Moreover, this positive impact of low doses of ethanol has been observed and confirmed by more recent studies. Indeed, Bhattacharya *et al*. (1985) showed that ethanol increased the root formation in mung bean, with a maximum reached at 0.2% (equivalent to 34.25 mM), and Wu *et a*l. (2019) reported that ethanol improved both biomass (at 0.0125 - 0.05 mL.L^-1^, equivalent to 0.21 - 0.86 mM) and micronutrient accumulation in oilseed rape. The characteristic response of hormesis can be observed in the results of transpiration of Gamay cuttings subjected to water stress and EtOH priming, in a companion article published in this issue (Ait Kaci *et al*., 2024). And finally, the curve shapes of cellular leakage observed in our study (Fig. 4), on two different plant species, strengthen the previous results. Thus, our results suggest that ethanol at physiological concentrations interacts with grapevine and *Arabidopsis* biological systems in a complex manner, where small amounts can activate adaptive and beneficial responses, while larger amounts overwhelm these systems, leading to toxicity. Given its potential role in activating adaptive responses, we hypothesise that ethanol may serve as a signalling molecule in plants and could be a potential messenger of stressful conditions.

Further research is needed to explore ethanol-induced hormesis in plants, specifically to elucidate how stimulation and inhibition vary with ethanol concentration. While the inhibition observed at high concentrations is expected as it aligns with the onset of toxicity, the stimulation effect at low concentrations warrants more detailed investigation. Considering ethanol status as a small molecule with significant diffusive capabilities, notably through membranes, and considering its hydrophobic nature, the manner by which cells perceive small variation in ethanol concentration is still a matter of research and debate (Diot *et al*., 2024).

## Conclusion

Our study demonstrated that grapevine cells sense ethanol at a physiological concentration, inducing a strong and rapid transcriptomic response, typical of stressors. This transcriptomic response had physiological repercussions, as evidenced by the thermotolerance biphasic dose-response curves, suggesting that ethanol could act as a potential molecular messenger in stressful situations in plant systems, rather than just being an end-product metabolite of hypoxic metabolisms. Harnessing this response could be a promising strategy to prepare plants for more challenging stress conditions. Indeed, plants biosynthesize various compounds, such as phytohormones and other metabolites, to adapt to adverse environments. Recent studies have demonstrated that exogenous treatment of plants with chemical compounds can enhance abiotic stress tolerance by inducing molecular and physiological defence mechanisms, a process known as chemical priming (Sako *et al*., 2020). The development of effective strategies that mitigate abiotic stress is essential for sustainable agriculture and food security, especially with continuing global population growth. Chemical priming is believed to represent a promising strategy for mitigating abiotic stress in crop plants. Given its low cost and environmental non-persistence, we propose that ethanol could be a good candidate for plant chemical priming and a valuable tool in agriculture, especially for mitigating environmental stress impacts on crops.

## Supplementary data

The following supplementary data are available at the bottom of this file or can be requested from the corresponding author, when dealing with Excel files.

### List of Supplementary Figures

**Fig. S1.** Heat-stress monitoring in Gamay cell cultures.

**Fig. S2.** Average weight of a callus of Vitis vinifera cv. Gamay as a function of development time after sub-culturing.

**Fig. S3.** Dissolved O_2_ profiles of Vitis vinifera L. cv. Gamay Fréaux cell cultures as a function of development time.

**Fig. S4.** Photographs of calli seen from above and photographs of calli cross-sections.

**Fig. S5.** Transcriptional dynamics showing the number of DEGs upregulated and downregulated, 6 and 24 hours after a 1 mM ethanol treatment, compared to the controls.

**Fig. S6.** Details of the 16 DEGs within the enriched GO term “Secondary metabolism process”.

**Fig. S7A and S7B.** Monitoring anthocyanin concentration and total polyphenol index of Gamay cell cultures after an initial 1 mM ethanol treatment or control treatment.

### List of Supplementary Tables

**Table S1.** Composition of the Gamay cell culture growth medium

**Table S2.** Table showing the ‘concentration-effect’ of the ethanol treatment in 3-week-old calli, at the time of treatment with the exogenous ethanol solution.

**Table S3A and S3B.** Quality of the alignment and Assignment rate to features (RNA-seq data)

**Table S4.** Results of the Differential Expression Analysis of the comparison 0 vs 1 mM EtOH at 6h

**Tables S5A and S5B.** Lists of the 386 functionally annotated DEGs for the 6h comparison, 1 mM versus 0mM: 254 Down-Regulated DEGs (A) and 132 Up-Regulated DEGs (B)

**Table S6.** Overall GO enrichment analysis results for the up-regulated DEG list (132 genes).

**Table S7.** Overall GO enrichment analysis results for the down-regulated DEG list (254 genes).

**Table S8.** Details of the up-regulated DEGs within the enriched GO term “Response to heat”, resulting from the GO enrichment analysis.

**Table S9.** Details of the down-regulated DEGs within the enriched GO term “Secondary metabolism process”, resulting from the GO enrichment analysis.

## Abbreviations

DEGs: Differentially Expressed Genes
EtOH: Ethanol
GO: Gene Ontology
HS: Heat Shock
HSP: Heat-Shock Protein
PCA: Principal Component Analysis
RNA-seq: RNA sequencing
sHSP: small Heat-Shock Protein

## Acknowledgments

We thank EUR TULIP and the Occitanie Region for the PhD grant to A. Diot. We also thank P. Thuleau for providing the *Arabidopsis thaliana* Col-0 seeds, L. Lemonnier and D. Saint-Martin for help in cell and plant cultures. T. Gillet for his help with oxygen measurements, and R. Althiab-Almasaud and P. Fraysse for their help with RNA-seq sample preparation. Then we want to thank M. Bouzayen and J. Pirrello for GBF team co-direction and side-funding. Finally, we are grateful to the GenoToul bioinformatics platform in Toulouse (Bioinfo Genotoul, https://bioinfo.genotoul.fr/) for providing computing resources.

## Author contributions

C.C. and A.Diot conceptualised the project and designed the research. A.Diot performed the experimentations. G.M. and A.Djari performed the RNA-seq data cleaning, reads alignment and quality assessment of the reads. A.Diot, E.M. and G.M. performed the analyses on R and O.D.V., G.M. and A.Diot made the R graphs. A.Diot and C.C. wrote the original manuscript draft, and all co-authors contributed to editing the manuscript. A.Diot, S.B. and C.C. were involved in funding acquisition, C.C. and S.B. were involved in A.Diot’s PhD supervision.

## Conflict of interest

The authors declare they have no conflict of interest.

## Funding

This study was supported by the École Universitaire de Recherche TULIP-GS (ANR-18-EURE-0019), which provided half of a PhD grant to A. Diot and a full PhD grant to Olivia D.V. All authors are also grateful to the Occitanie Region for funding the other half of the PhD grant to A. Diot. The research was also partly funded by the VitiFunGen project supported by the Fondation Jean Poupelain (Cognac, France), the Labex TULIP (ANR-10-LABX-41) and by OxyFruit ANR (ANR-23-CE20-0001). Additionally, we thank the Université Paul Sabatier, CNRS, and Toulouse INP for their contributions to our research.

## Data availability

The data underlying this article will be shared upon request to the corresponding author.

## Companion article

This article is one of two companion articles submitted to the same journal. The second article by Ait Kaci et al. is entitled “Ethanol reduces grapevine water consumption by limiting transpiration”.

---

**Supplementary Fig. S1.**
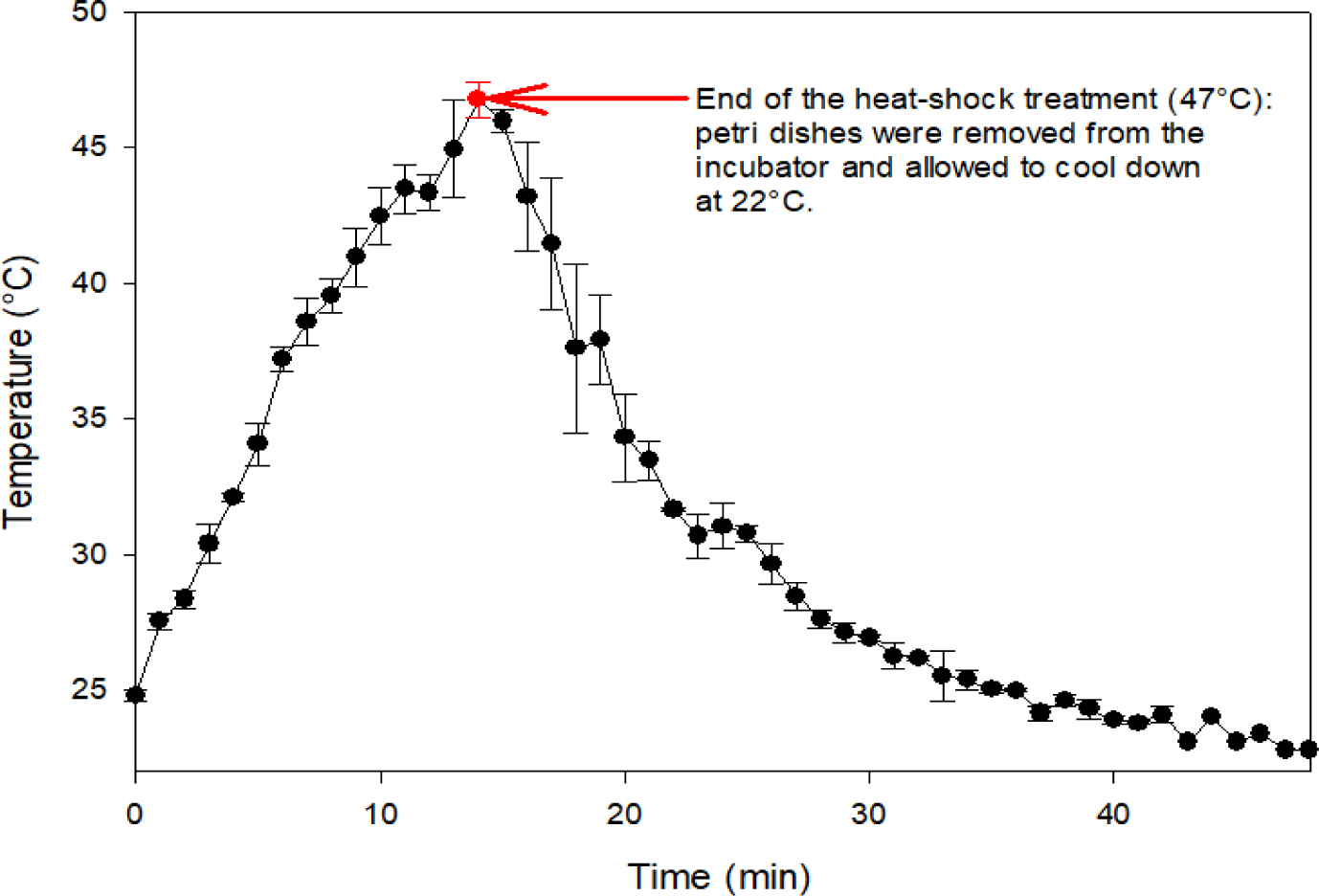
Heat-stress monitoring in Gamay cell cultures. The heat stress was applied in an incubator under dark conditions. The data represent the average temperature ± SD at the surface of the growth medium (n=3).

**Supplementary Fig. S2.**
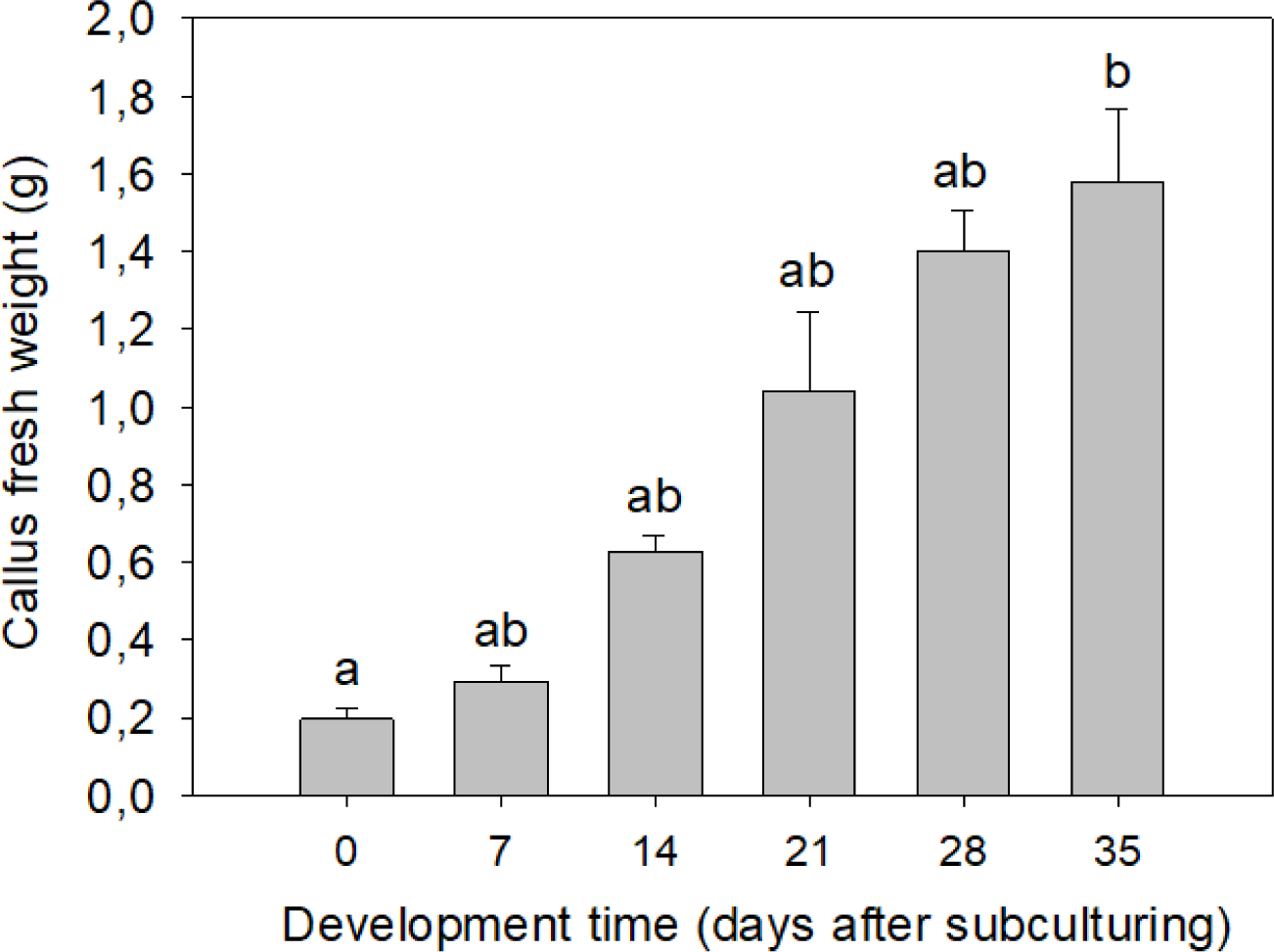
Average weight of a callus of *Vitis vinifera* cv. Gamay as a function of development time after sub-culturing. n = 12 different calli, error bars show SD, different small letters indicate significant differences at 0.05 (Dunn’s test).

**Supplementary Fig. S3.**
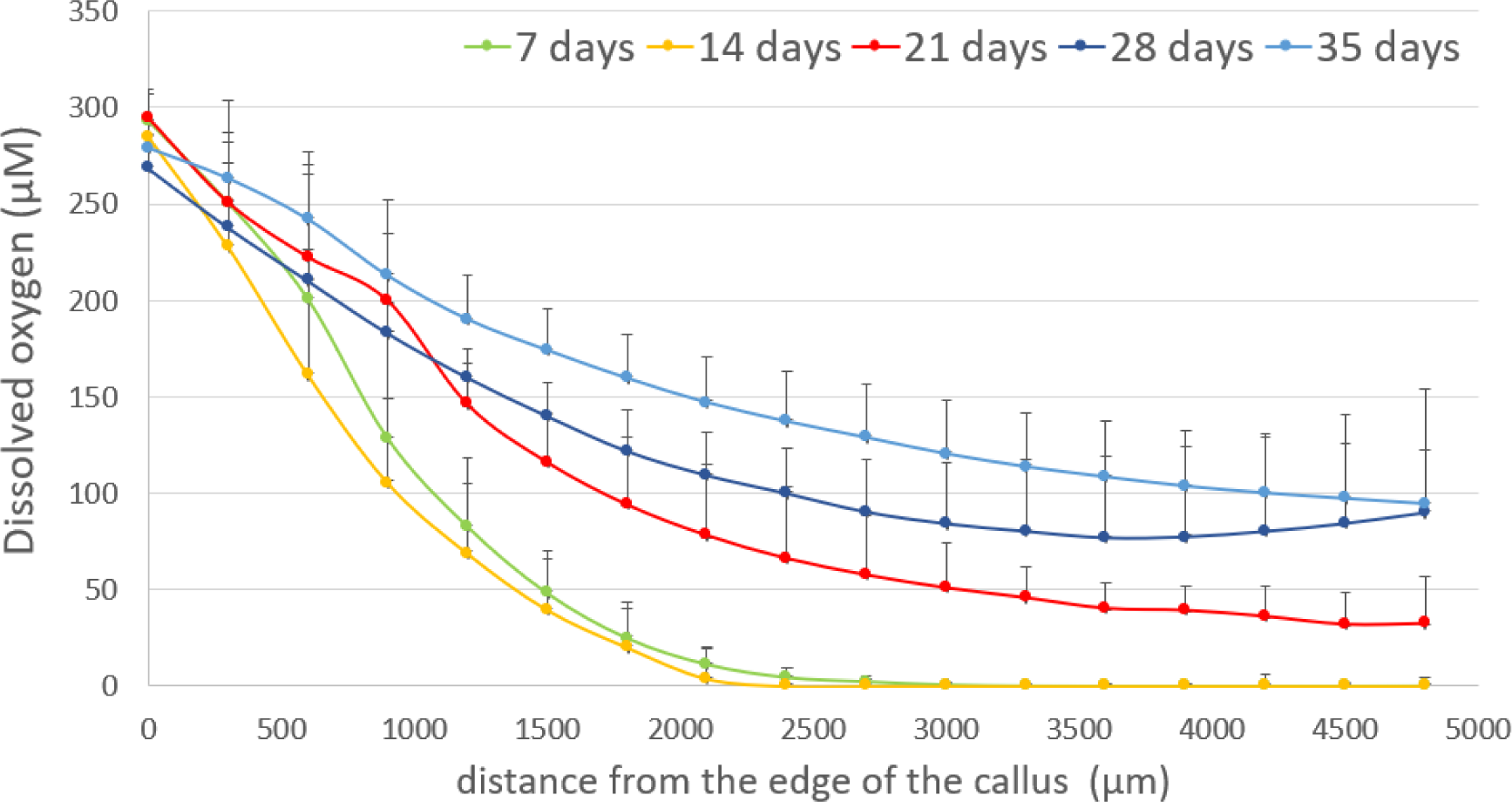
Dissolved O_2_ profiles of *Vitis vinifera* L. cv. Gamay Fréaux cell cultures as a function distance from the edge of the callus (in µm) and the development time (7, 14, 21, 28 and 35 days), n = 4 different calli, error bars show SD.

**Supplementary Fig. S4.**
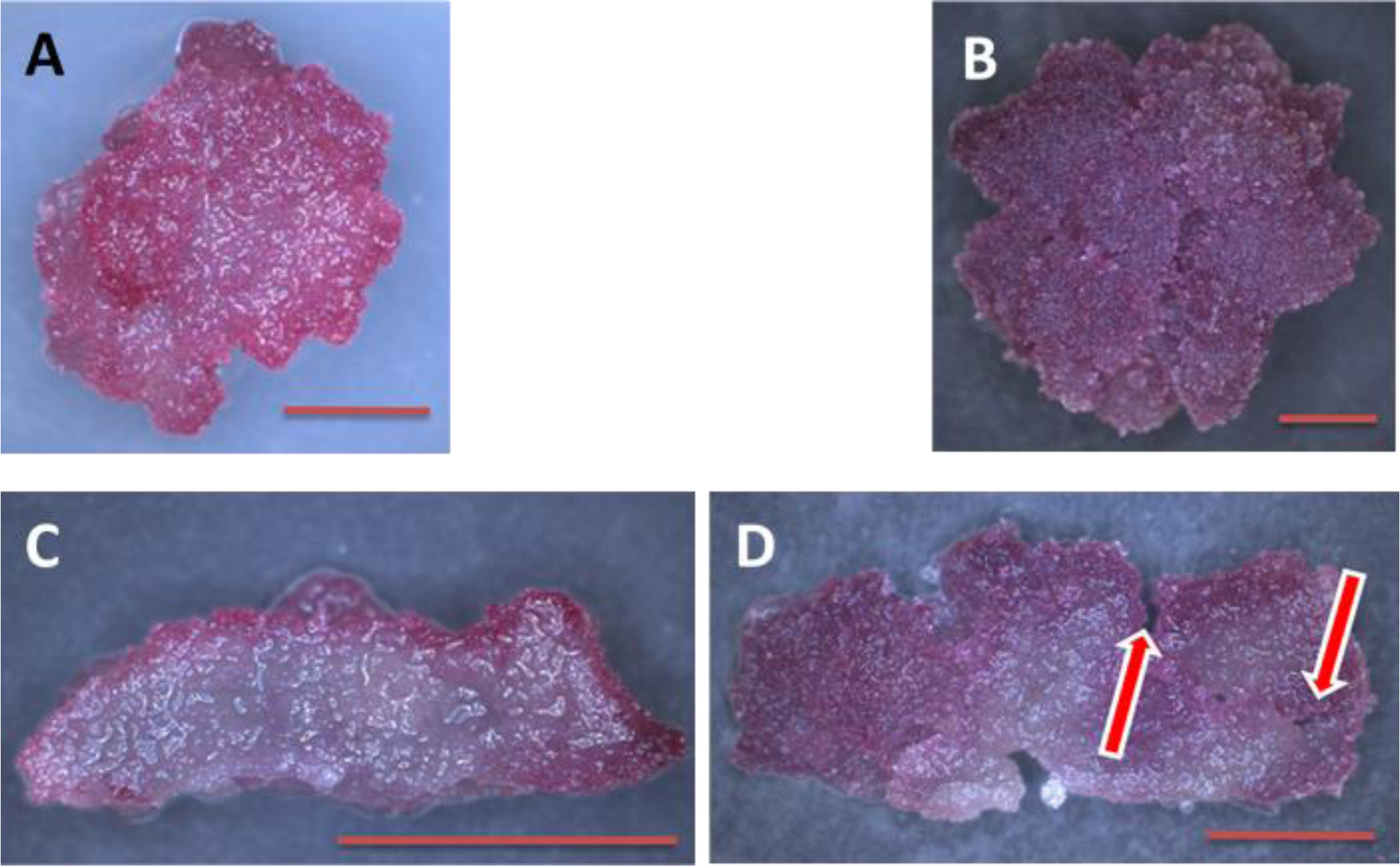
Photographs of calli seen from above (**A** and **B**) and photographs of calli cross-sections (**C** and **D**). Calli are either 14 days (**A** and **C**) or 35 days (**B** and **D**) old for comparison. Scale: the orange bar represents 0.5 cm. Red arrows show invaginations and gaseous interstices.

**Supplementary Fig. S5.**
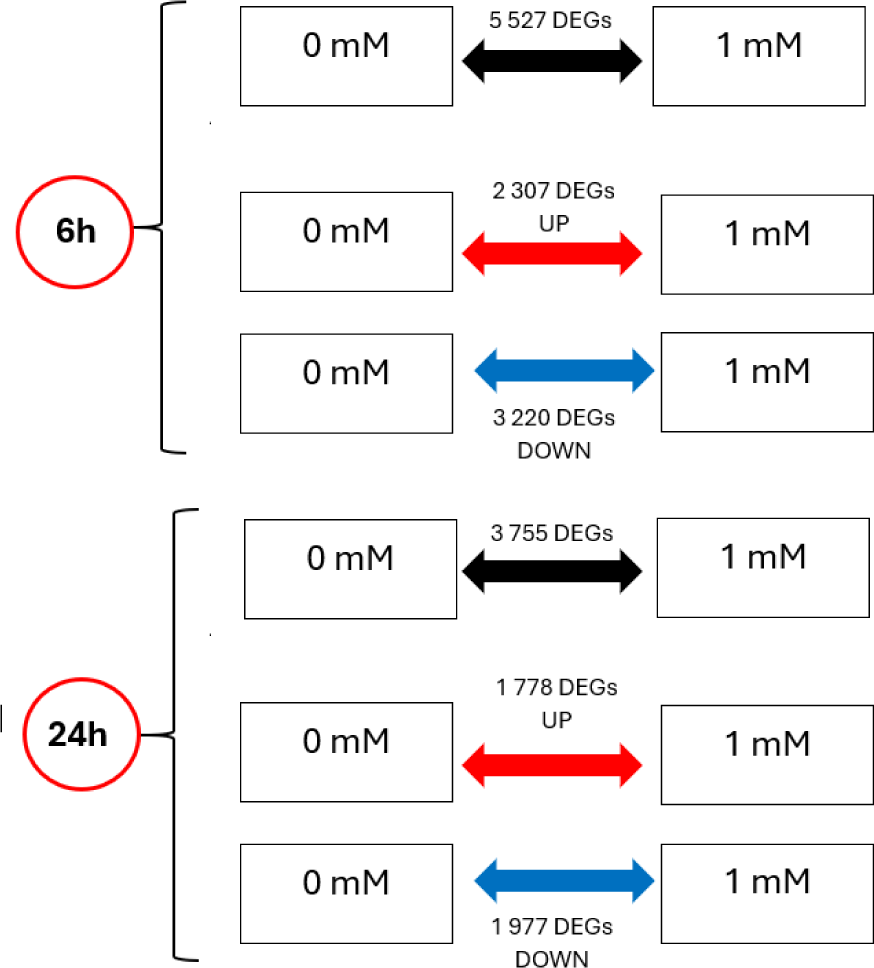
Transcriptional dynamics showing the number of DEGs upregulated and downregulated, 6 and 24 hours after a 1 mM ethanol treatment, compared to the controls. DEGs shown in this figure are all the genes with a p-adj < 0.05, with no regard to their Log2FoldChange.

**Supplementary Fig. S6.**
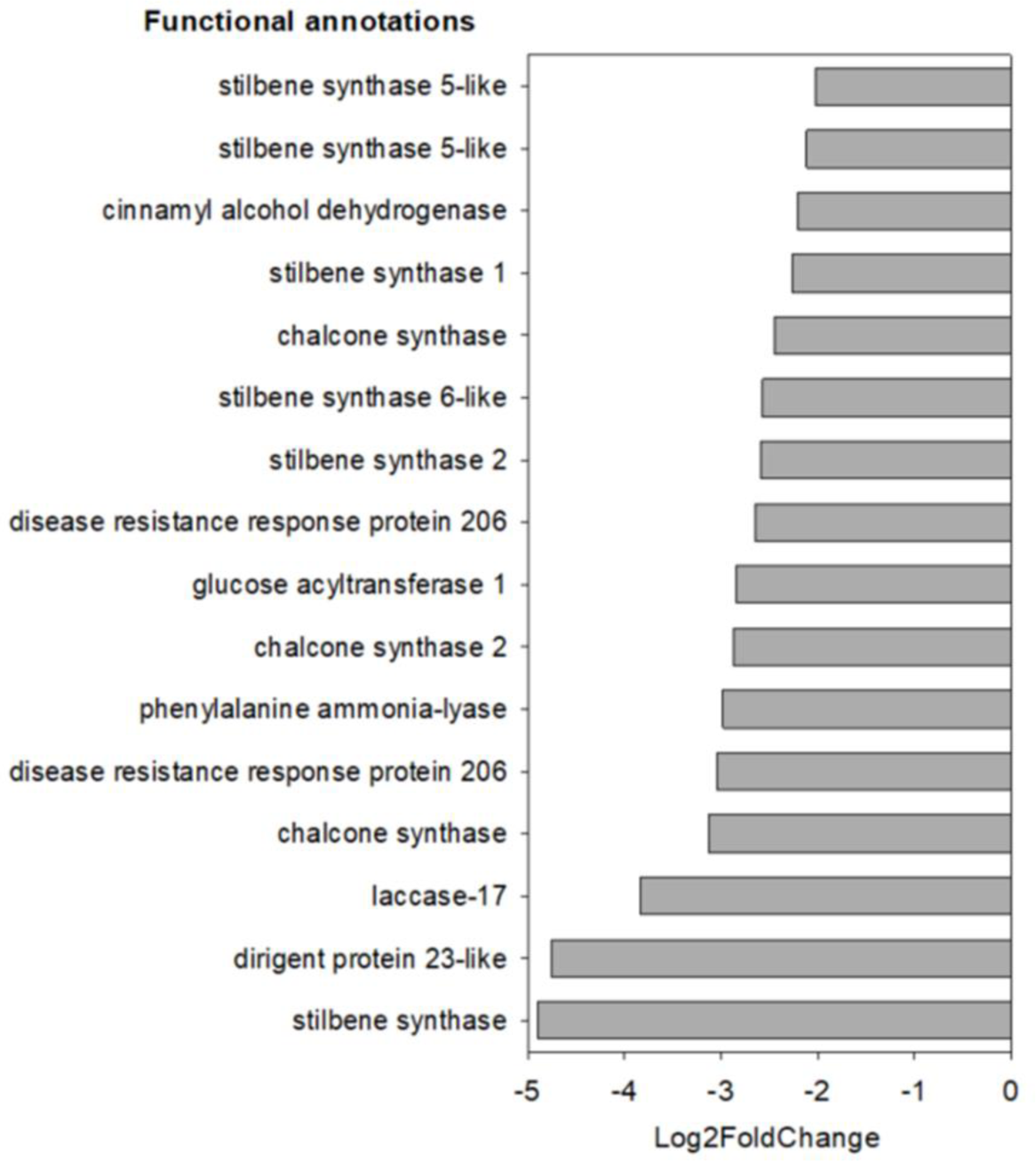
Details of the 16 DEGs, within the enriched GO term “secondary metabolism process”, ordered by their respective fold changes, comparing 1 mM ethanol treatment to control, at 6 hour post-treatment.

**Supplementary Fig. S7:**
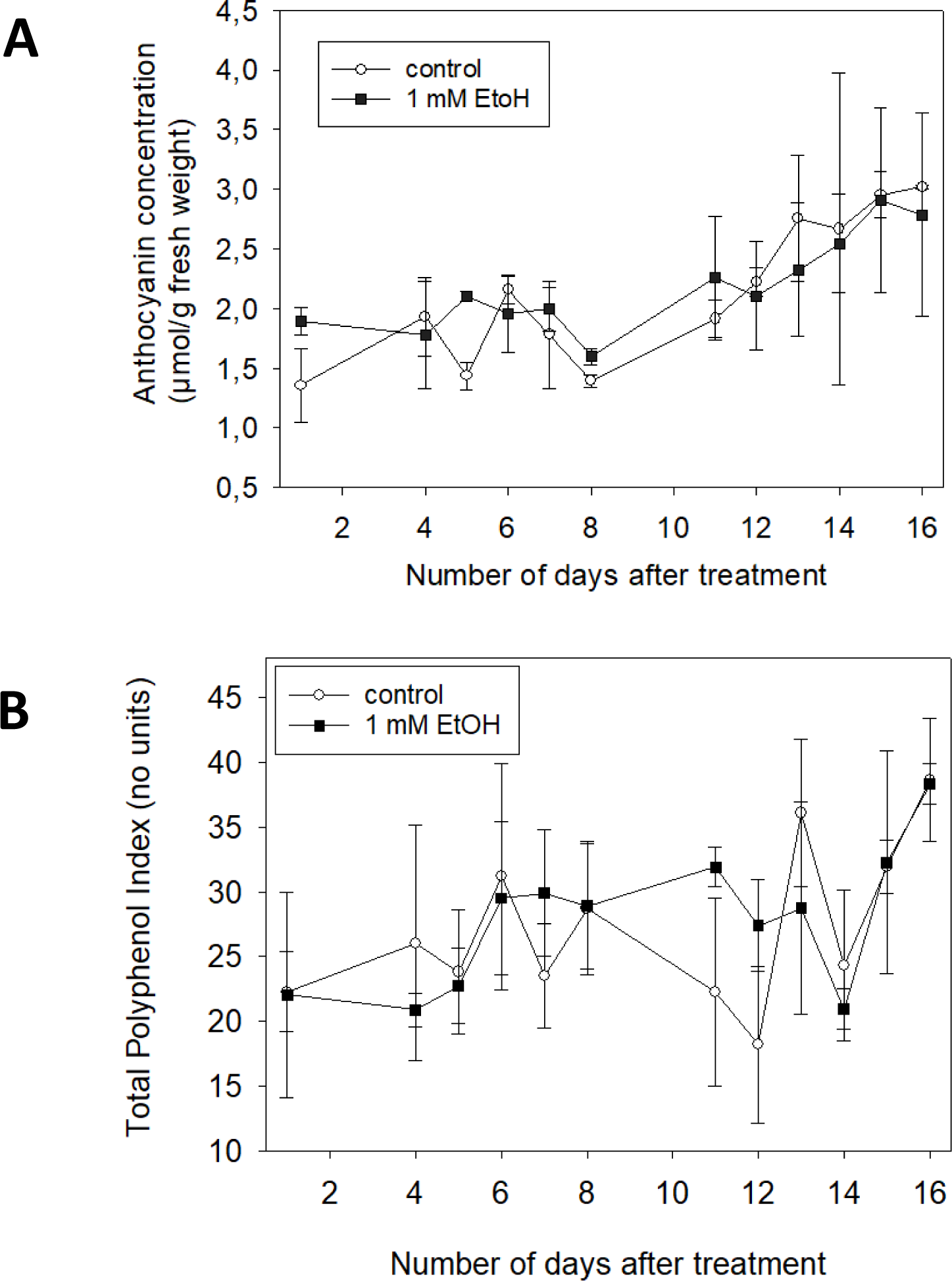
Temporal changes in **A)** anthocyanin concentration and **B)** in the total polyphenol index (TPI) in grapevine cell cultures following an initial treatment of 1 mM ethanol (black squares) or a control treatment (white circles). Data represent the mean ± SD of three samples.

**Supplementary Table S1.** Composition of growth medium for *Vitis vinifera* Gamay Fréaux cell cultures *(as it is not easy to find in old articles)*

| Macronutrients | mg/L |
| --- | --- |
| KNO<sub>3</sub> | 2,500 |
| (NH<sub>4</sub>)<sub>2</sub>SO<sub>4</sub> | 150 |
| CaCl<sub>2</sub>, 2H<sub>2</sub>O | 150 |
| NaH<sub>2</sub>PO<sub>4</sub>, 2H<sub>2</sub>O | 150 |
| MgSO<sub>4</sub>, 7H<sub>2</sub>O | 250 |
| Microelements | mg/L |
| MnSO<sub>4</sub>, H<sub>2</sub>O | 16.9 |
| ZnSO<sub>4</sub>, 7H<sub>2</sub>O | 8.6 |
| H<sub>3</sub>BO<sub>3</sub> | 6.2 |
| KI | 0.83 |
| NaMoO<sub>4</sub>, 2H<sub>2</sub>O | 0.25 |
| Cu SO<sub>4</sub>, 5H<sub>2</sub>O | 0.025 |
| CoCl<sub>2</sub>, 6H<sub>2</sub>O | 0.025 |
| Iron/EDTA | mg/L |
| FeSO<sub>4</sub>, 7H<sub>2</sub>O | 27.85 |
| Na<sub>2</sub>EDTA | 37.85 |
| Vitamins | mg/L |
| Myoinositol | 100 |
| D-pantothenic acid, hemicalcium salt | 1 |
| Biotin | 0.01 |
| Nicotinic acid | 1 |
| Thiamine-HCl | 1 |
| Pyridoxine-HCl | 1 |
| Organic compounds | mg/L |
| Sucrose | 20,000 |
| Casein hydrolate | 250 |
| Growth regulators | mg/L |
| Kinetin | 0.2 |
| 1-naphthaleneacetic acid | 0.1 |
| Agar | 9 g/L |
| NaOH | Adjust to pH = 6 |

**Supplementary Table S2.**
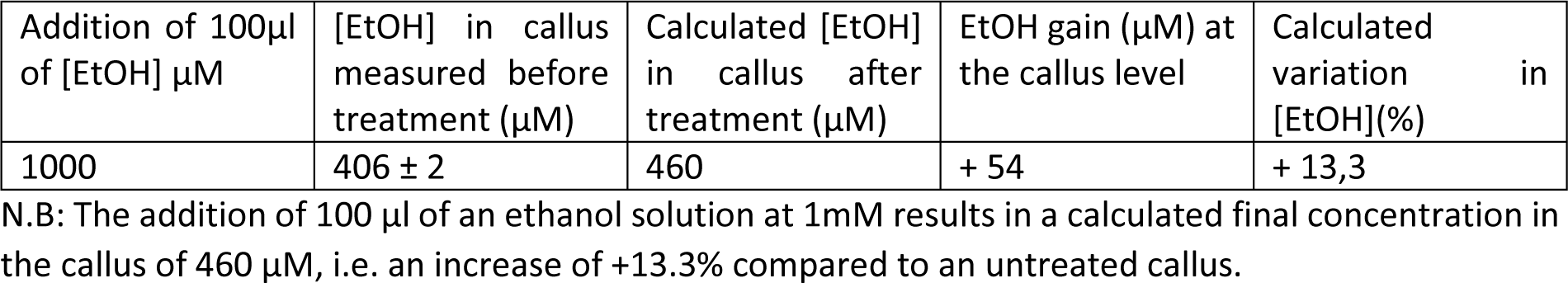
Table showing the ‘concentration-effect’ of the ethanol treatment in 3-week-old calli, at the time of treatment with the exogenous ethanol solution. The endogenous ethanol concentration was measured in 3 calli. The final ethanol concentration after treatment was calculated to account for the light ethanol change generated by the 1 mM EtOH treatment.

**Supplementary Table S3A.** Quality of the alignment - Percentage of uniquely mapped reads on the *Vitis vinifera* cv. Chasselas reference genome (Djari *et al*., 2024).

| Treatment duration | sample designation | treatment | % of uniquely mapped reads |
| --- | --- | --- | --- |
| 6h | etha_6h_0mM_1 | control | 93.12 |
| 6h | etha_6h_0mM_2 | control | 93.14 |
| 6h | etha_6h_0mM_3 | control | 92.48 |
| 6h | etha_6h_1mM_2 | Ethanol 1 mM | 93.51 |
| 6h | etha_6h_1mM_3 | Ethanol 1 mM | 92.89 |
| 6h | etha_6h_1mM_4 | Ethanol 1 mM | 93.45 |
| 24h | etha_24h_0µM_2 | control | 92.55 |
| 24h | etha_24h_0µM_3 | control | 92.48 |
| 24h | etha_24h_0µM_4 | control | 92.09 |
| 24h | etha_24h_1000µM_1 | Ethanol 1 mM | 92.44 |
| 24h | etha_24h_1000µM_2 | Ethanol 1 mM | 92.96 |
| 24h | etha_24h_1000µM_3 | Ethanol 1 mM | 92.84 |

**Supplementary Table S3B.** Assignment rate to features on the *Vitis vinifera* cv. Chasselas reference genome (Djari *et al*., 2024).

| Treatment duration | sample designation | treatment | % of reads assigned to features |
| --- | --- | --- | --- |
| 6h | etha_6h_0mM_1 | control | 83.40 |
| 6h | etha_6h_0mM_2 | control | 83.07 |
| 6h | etha_6h_0mM_3 | control | 81.77 |
| 6h | etha_6h_1mM_2 | Ethanol 1 mM | 83.01 |
| 6h | etha_6h_1mM_3 | Ethanol 1 mM | 82.11 |
| 6h | etha_6h_1mM_4 | Ethanol 1 mM | 82.95 |
| 24h | etha_24h_0µM_2 | control | 80.69 |
| 24h | etha_24h_0µM_3 | control | 82.45 |
| 24h | etha_24h_0µM_4 | control | 80.93 |
| 24h | etha_24h_1000µM_1 | Ethanol 1 mM | 82.08 |
| 24h | etha_24h_1000µM_2 | Ethanol 1 mM | 83.56 |
| 24h | etha_24h_1000µM_3 | Ethanol 1 mM | 81.12 |

*Supplementary Tables S4 to S9 (Excel files) available on the publisher website*

## Notes

### Competing Interest Statement

The authors have declared no competing interest.

### Summary of Updates

We now mention all the funders, while some were not mentioned in a first version. Additionally, we have removed mention of the journal in which this manuscript was submitted. Supplementary files were added to the PDF. Finally, author names were rearranged.

## References

Abbas M, Sharma G, Dambire, C, Marquez J, Alonso-Blanco C, Proaño K, Holdsworth MJ (2022) An oxygen-sensing mechanism for angiosperm adaptation to altitude. Nature 606:565–569. 10.1038/s41586-022-04740-y

Agathokleous E, Kitao M, Calabrese EJ (2020) Hormesis: Highly generalizable and beyond laboratory. Trends Plant Sci 25:1076–1086. doi: 10.1016/j.tplants.2020.05.006

Aghdam MS, Sevillano L, Flores FB, Bodbodak S (2013) Heat shock proteins as biochemical markers for postharvest chilling stress in fruits and vegetables. Sci Hort 160:54–64. 10.1016/j.scienta.2013.05.020

Ait Kaci N, Quinquiry B, Diot A, Yobregat O, Pellegrino A, Maury P, Chervin C (2024) Ethanol reduces grapevine water consumption by limiting transpiration. Submitted (companion paper).

Alkorta I, Epelde L, Garbisu C (2017) Environmental parameters altered by climate change affect the activity of soil microorganisms involved in bioremediation. FEMS 364:fnx200. 10.1093/femsle/fnx200

Althiab-Almasaud R, Chen Y, Maza E, Djari A, Frasse P, Mollet JC, Mazars C, Jamet E, Chervin C (2021) Ethylene signaling modulates tomato pollen tube growth through modifications of cell wall remodeling and calcium gradient. Plant J 107:893–908. 10.1111/tpj.15353

Altschuler M, Mascarenhas JP (1982) Heat shock proteins and effects of heat shock in plants. Plant Mol Biol 1:103–115. 10.1007/BF00024974

Baird NA, Turnbull DW, Johnson EA (2006) Induction of the heat shock pathway during hypoxia requires regulation of heat shock factor by hypoxia-inducible factor-1. J Biol Chem 281:38675–38681. 10.1074/jbc.M608013200

Basha E, Jones C, Wysocki V, Vierling E (2010) Mechanistic differences between two conserved classes of small heat shock proteins found in the plant cytosol. J Biol Chem 285:11489–11497. 10.1074/jbc.M109.074088

Bhattacharya S, Bhattacharya NC, Bhatnagar VB (1985) Effect of ethanol, methanol and acetone on rooting etiolated cuttings of Vigna radiata in presence of sucrose and auxin. Ann Bot 55:143–145. 10.1093/oxfordjournals.aob.a086885

Bashir K, Todaka D, Rasheed S, Matsui A, Ahmad Z, Sako K, Utsumi Y, Vu AT, Tanaka M, Takahashi S, Ishida J (2022) Ethanol-mediated novel survival strategy against drought stress in plants. Plant Cell Physiol 63:1181–1192. 10.1093/pcp/pcac114

Bilgin DD, Zavala JA, Zhu JIN, Clough SJ, Ort DR, DeLucia EH (2010) Biotic stress globally downregulates photosynthesis genes. Plant Cell Environ 33:1597–1613. 10.1111/j.1365-3040.2010.02167.x

Borisjuk L, Rolletschek H (2009) The oxygen status of the developing seed. New Phytol 182:17–30. 10.1111/j.1469-8137.2008.02752.x

Brazel AJ, Graciet E (2023) Complexity of abiotic stress stimuli: Mimicking hypoxic conditions experimentally on the basis of naturally occurring environments. In Plant Abiotic Stress Signaling. New York, Springer US, pp 23–48. 10.1007/978-1-0716-3044-0_2

Bui LT, Novi G, Lombardi L, Iannuzzi C, Rossi J, Santaniello A, Mensuali A, Corbineau F, Giuntoli B, Perata P, Zaffagnini M (2019) Conservation of ethanol fermentation and its regulation in land plants. J Exp Bot 70:1815–1827. 10.1093/jxb/erz052

Calabrese EJ, Baldwin LA (2002) Defining hormesis. Human & Experimental Toxicology 21(2), 91–97. 10.1191/0960327102ht217oa

Calabrese EJ, Baldwin LA (2003) Ethanol and hormesis. Critical Reviews in Toxicology 33 407–424. 10.1080/713611043

Canaguier A, Grimplet J, Di Gaspero G, Scalabrin S, Duchêne E, Choisne N, Mohellibi N, Guichard C, Rombauts S, Le Clainche I, Berard A (2017) A new version of the grapevine reference genome assembly (12X. v2) and of its annotation (VCost. v3). Genom data 14:56. 10.1016/j.gdata.2017.09.002

Cetó X, Gutiérrez JM, Gutiérrez M, Céspedes F, Capdevila J, Mínguez S, Jiménez-Jorquera C, Del Valle M (2012) Determination of total polyphenol index in wines employing a voltametric electronic tongue. Anal Chim Acta 732:172–179. 10.1016/j.aca.2012.02.026

Chapra SC, Camacho LA, McBride GB (2021) Impact of global warming on dissolved oxygen and BOD assimilative capacity of the world’s rivers: modeling analysis. Water 13:2408. 10.3390/w13172408

Chen Y, Chen X, Wang H, Bao Y, Zhang W (2014) Examination of the Leaf Proteome During Flooding Stress and the Induction of Programmed Cell Death in Maize. Proteome Sci 12:33. 10.1186/1477-5956-12-33

Chervin C, Elkereamy A, Roustan JP, Faragher JD, Latché A, Pech JC, Bouzayen M (2001) An ethanol spray at veraison enhances colour in red wines. Aust J Grape Wine Res 7:144–145. 10.1111/j.1755-0238.2001.tb00202.x

Chirinos X, Ying S, Rodrigues MA, Maza E, Djari A, Hu G, Liu M, Purgatto E, Fournier S, Regad F, Bouzayen M (2023) Transition to ripening in tomato requires hormone-controlled genetic reprogramming initiated in gel tissue. Plant Physiol 191:610–625. 10.1093/plphys/kiac464

Choi MR, Jung KH, Park JH, Das ND, Chung MK, Choi IG, Lee BC, Park KS, Chai YG (2011) Ethanol-induced small heat shock protein genes in the differentiation of mouse embryonic neural stem cells. Arch Toxicol 85:293–304. 10.1007/s00204-010-0591-z

Daniel K, Hartman S (2024) How plant roots respond to waterlogging. J Exp Bot 75:511–525. 10.1093/jxb/erad332

Das AK, Anik TR, Rahman MM, Keya SS, Islam MR, Rahman MA, Sultana S, Ghosh PK, Khan S, Ahamed T, Ghosh TK (2022) Ethanol treatment enhances physiological and biochemical responses to mitigate saline toxicity in soybean. Plants 11: 272. 10.3390/plants11030272

Diot A, Groth G, Blanchet S, Chervin C (2024) Responses of animals and plants to physiological doses of ethanol: a molecular messenger of hypoxia? FEBS J 291:1102–1110. 10.1111/febs.17056

Djari A, Madignier G, Di Valentin O, Gillet T, Frasse P, Djouhri A, Hu G, Julliard S, Liu M, Zhang Y, Regad F (2024) Haplotype-resolved genome assembly and implementation of VitExpress, an open interactive transcriptomic platform for grapevine. Proc Natl Acad Sci USA 121:e2403750121. 10.1073/pnas.2403750121

Dukowic-Schulze S, van der Linde K (2021) Oxygen, secreted proteins and small RNAs: mobile elements that govern anther development. Plant Reprod 34:1–19. 10.1007/s00497-020-00401-0

Edgar R, Domrachev M, Lash AE (2002) Gene Expression Omnibus: NCBI gene expression and hybridization array data repository. Nucleic Acid Res 30:207–210. 10.1093/nar/30.1.207

Erofeeva EA (2022) Hormesis in plants: Its common occurrence across stresses. Curr Op Toxicol 30:100333. 10.1016/j.cotox.2022.02.006

Geigenberger P, Fernie AR, Gibon Y, Christ M, Stitt M (2000) Metabolic activity decreases as an adaptive response to low internal oxygen in growing potato tubers. Biol Chem 381:723–40. 10.1515/BC.2000.093

Geigenberger P (2003) Response of plant metabolism to too little oxygen. Curr Op Plant Biol 6:247–256. 10.1016/s1369-5266(03)00038-4

Guan JC, Jinn TL, Yeh CH, Feng SP, Chen YM, Lin CY (2004) Characterization of the genomic structures and selective expression profiles of nine class I small heat shock protein genes clustered on two chromosomes in rice (Oryza sativa L.). Plant Mol Biol 56:795–809. 10.1007/s11103-004-5182-z

Guan L, Dai Z, Wu BH, Wu J, Merlin I, Hilbert G, Renaud C, Gomès E, Edwards E, Li SH, Delrot S (2016) Anthocyanin biosynthesis is differentially regulated by light in the skin and flesh of white-fleshed and teinturier grape berries. Planta 243:23–41. 10.1007/s00425-015-2391-4

Haslbeck M, Vierling E (2015) A first line of stress defense: small heat shock proteins and their function in protein homeostasis. J Mol Biol 427:1537–1548. 10.1016/j.jmb.2015.02.002

Hellwig S, Drossard J, Twyman RM, Fischer R (2004) Plant cell cultures for the production of recombinant proteins. Nature Biotech 22:1415–1422. 10.1038/nbt1027

Hu C, Yang J, Qi Z, Wu H, Wang B, Zou F, Mei H, Liu J, Wang W, Liu Q (2022) Heat shock proteins: Biological functions, pathological roles, and therapeutic opportunities. MedComm 3(3), e161. 10.1002/mco2.161

Ilík P, Špundová M, Šicner M, Melkovičová H, Kučerová Z, Krchňák P, Fürst T, Večeřová K, Panzarová K, Benediktyová Z, Trtílek M (2018) Estimating heat tolerance of plants by ion leakage: a new method based on gradual heating. New Phytol 218:1278–1287. 10.1111/nph.15097

Janowska MK, Baughman HE, Woods CN, Klevit RE (2019) Mechanisms of small heat shock proteins. Cold Spring Harb Perspec Biol 11:a034025. 10.1101/cshperspect.a034025

Jethva J, Schmidt RR, Sauter M, Selinski J (2022) Try or die: Dynamics of plant respiration and how to survive low oxygen conditions. Plants 11:205. 10.3390/plants11020205

Jiang X, Feng K, Yang X (2015) In vitro antifungal activity and mechanism of action of tea polyphenols and tea saponin against *Rhizopus stolonifer*. J Mol Microbiol Biotech 25:269–276. 10.1159/000430866

Kimmerer TW, Kozlowski TT (1982) Ethylene, ethane, acetaldehyde, and ethanol production by plants under stress. Plant Physiol 69:840–847. 10.1104/pp.84.4.1204

Kong J, Wu J, Guan L, Hilbert G, Delrot S, Fan P, Liang Z, Wu B, Matus JT, Gomès E, Dai Z (2021) Metabolite analysis reveals distinct spatio-temporal accumulation of anthocyanins in two teinturier variants of cv.‘Gamay’grapevines (Vitis vinifera L.). Planta 253:1–18. 10.1007/s00425-021-03613-4

Kuo HF, Tsai YF, Young LS, Lin CY. (2000) Ethanol treatment triggers a heat shock-like response but no thermotolerance in soybean (Glycine max cv. Kaohsiung No. 8) seedlings. Plant Cell Envir 23:1099-1108. 10.1046/j.1365-3040.2000.00621.x

Lambri M, Torchio F, Colangelo D, Segade SR, Giacosa S, De Faveri DM, Gerbi V, Rolle L (2015) Influence of different berry thermal treatment conditions, grape anthocyanin profile, and skin hardness on the extraction of anthocyanin compounds in the colored grape juice production. Food Res Int 77:584–590. 10.1016/j.foodres.2015.08.027

Lee GJ, Pokala N, Vierling E (1995) Structure and in vitro molecular chaperone activity of cytosolic small heat shock proteins from pea. J Biol Chem 270:10432–10438. 10.1074/jbc.270.18.10432

Lee GJ, Roseman AM, Saibil HR, Vierling E (1997) A small heat shock protein stably binds heat-denatured model substrates and can maintain a substrate in a folding-competent state. EMBO J 16:659–671. 10.1093/emboj/16.3.659

Li GC (1983) Induction of thermotolerance and enhanced heat shock protein synthesis in Chinese hamster fibroblasts by sodium arsenite and by ethanol. J Cell Physiol 115:116–122. 10.1002/jcp.1041150203

Liu GT, Wang JF, Cramer G, Dai ZW, Duan W, Xu HG, Wu BH, Fan PG, Wang LJ, Li SH (2012) Transcriptomic analysis of grape (Vitis vinifera L.) leaves during and after recovery from heat stress. BMC Plant Biol 12:1–10. 10.1186/1471-2229-12-174

Lindquist S, Craig EA (1988) The heat-shock proteins. Ann Rev Genet 22:631–677. 10.1146/annurev.ge.22.120188.003215

Loreti E, Perata P (2020) The many facets of hypoxia in plants. Plants 9: 745. 10.3390/plants9060745

Love MI, Huber W, Anders S (2014) Moderated estimation of fold change and dispersion for RNA-seq data with DESeq2. Genome Biol 15:1–21. 10.1186/s13059-014-0550-8

Maza E, Frasse P, Senin P, Bouzayen M, Zouine M (2013) Comparison of normalization methods for differential gene expression analysis in RNA-Seq experiments: a matter of relative size of studied transcriptomes. Commun Integr Biol 6:e25849. 10.4161/cib.25849

Matsui A, Todaka D, Tanaka M, Mizunashi K, Takahashi S, Sunaoshi Y, Tsuboi Y, Ishida J, Bashir K, Kikuchi J, Kusano M (2022) Ethanol induces heat tolerance in plants by stimulating unfolded protein response. Plant Mol Biol 110:131–145. 10.1007/s11103-022-01291-8

McLoughlin F, Basha E, Fowler ME, Kim M, Bordowitz J, Katiyar-Agarwal S, Vierling E (2016) Class I and II small heat shock proteins together with HSP101 protect protein translation factors during heat stress. Plant Physiol 172:1221–1236. 10.1104/pp.16.00536

Meitha K, Agudelo-Romero P, Signorelli S, Gibbs DJ, Considine JA, Foyer CH, Considine MJ (2018) Developmental control of hypoxia during bud burst in grapevine. Plant Cell Environ 41:1154–1170. 10.1111/pce.13141

Middleton W, Jarvis BC, Booth A (1978) The effects of ethanol on rooting and carbohydrate metabolism in stem cuttings of Phaseolus aureus Roxb. New Phytol 81:279–285. 10.1111/j.1469-8137.1978.tb02633.x

Park CJ, Seo YS. (2015) Heat shock proteins: a review of the molecular chaperones for plant immunity. Plant Pathol J 31:323. 10.5423/ppj.rw.08.2015.0150

Parsell DA, Lindquist S (1993) The function of heat-shock proteins in stress tolerance: degradation and reactivation of damaged proteins. Ann Rev Genet 27:437–496. 10.1146/annurev.ge.27.120193.002253

Patra M, Salonen E, Terama E, Vattulainen I, Faller R, Lee BW, Holopainen J, Karttunen M (2006) Under the influence of alcohol: the effect of ethanol and methanol on lipid bilayers. Biophys J 90:1121–1135. 10.1529/biophysj.105.062364

Pignataro L, Miller AN, Ma L, Midha S, Protiva P, Herrera DG, Harrison NL (2007) Alcohol regulates gene expression in neurons via activation of heat shock factor 1. J Neurosci 27:12957–12966. 10.1523/jneurosci.4142-07.2007

Plesset J, Palm C, McLaughlin CS (1982) Induction of heat shock proteins and thermotolerance by ethanol in Saccharomyces cerevisiae. Biochem Biophys Res Comm 108:1340–1345. 10.1016/0006-291X(82)92147-7

Raymond P, Saglio P, Ricard B (1995) Réponse au manque d’oxygène dans les tissus végétaux. Cahiers Agric 4:343–350. https://revues.cirad.fr/index.php/cahiers-agricultures/article/view/29908/29668

Rolletschek H, Borisjuk L, Koschorreck M, Wobus U, Weber H (2002) Legume embryos develop in a hypoxic environment. J Exp Bot 53:1099–1107. 10.1093/jexbot/53.371.1099

Scott BR (2008) Low-dose-radiation stimulated natural chemical and biological protection against lung cancer. Dose-Response 6:299–318. 10.2203/dose-response.07-025.scott

Savvides A, Ali S, Tester M, Fotopoulos V (2016) Chemical priming of plants against multiple abiotic stresses: mission possible? Trends Plant Sci 21:329–340. 10.1016/j.tplants.2015.11.003

Sako K, Nguyen HM, Seki M (2020) Advances in chemical priming to enhance abiotic stress tolerance in plants. Plant Cell Physiol 61: 1995–2003. 10.1093/pcp/pcaa119

Sato F (2013) Characterization of plant functions using cultured plant cells, and biotechnological applications. Biosci Biotechnol Biochem 77:1–9. 10.1271/bbb.120759

Schwacke R, Ponce-Soto GY, Krause K, Bolger AM, Arsova B, Hallab A, Gruden K, Stitt M, Bolger ME, Usadel B (2019) MapMan4: a refined protein classification and annotation framework applicable to multi-omics data analysis. Mol Plant 12:879–892. 10.1016/j.molp.2019.01.003

Smetanska I (2008) Production of Secondary Metabolites Using Plant Cell Cultures. Adv Biochem Eng Biotechnol 111:187–228. 10.1007/10_2008_103

Sun Y, MacRae TH (2005) Small heat shock proteins: molecular structure and chaperone function. Cell Mol Life Sci 62:2460–2476. 10.1007/s00018-005-5190-4

Todaka D, Quynh DTN, Tanaka M, Utsumi Y, Utsumi C, Ezoe A, Takahashi S, Ishida J, Kusano M, Kobayashi M, Saito K (2024) Application of ethanol alleviates heat damage to leaf growth and yield in tomato. Front Plant Sci 15:1325365. 10.3389/fpls.2024.1325365

Triantaphylides C, Nespoulous L, Chervin C 1993. Ammonium requirement for radiation-induced accumulation of polyamines in suspension-cultured grape cells. Physiol Plant 87:389–395. 10.1111/j.1399-3054.1993.tb01746.x

Tsvetkova NM, Horváth I, Török Z, Wolkers WF, Balogi Z, Shigapova N, Crowe LM, Tablin F, Vierling E, Crowe JH, Vigh L (2002) Small heat-shock proteins regulate membrane lipid polymorphism. Proc Natl Acad Sci USA. 99:13504–9. 10.1073/pnas.192468399

Ul Haq S, Khan A, Ali M, Khattak AM, Gai WX, Zhang HX, Wei AM, Gong ZH (2019) Heat shock proteins: dynamic biomolecules to counter plant biotic and abiotic stresses. Int J Mol Sci 20:5321. 10.3390/ijms20215321

Vierling E (1991) The roles of heat shock proteins in plants. Ann Rev Plant Phys Plant Mol Biol 42:579–620. 10.1146/annurev.pp.42.060191.003051

Vitrac X, Larronde F, Krisa S, Decendit A, Deffieux G, Mérillon JM (2000) Sugar sensing and Ca2+– calmodulin requirement in Vitis vinifera cells producing anthocyanins. Phytochem 53: 659–665. 10.1016/s0031-9422(99)00620-2

Wang W, Vinocur B, Shoseyov O, Altman A (2004) Role of plant heat-shock proteins and molecular chaperones in the abiotic stress response. Trends Plant Sci 9:244–252. 10.1016/j.tplants.2004.03.006

Wang B, Lin L, Yuan X, Zhu Y, Wang Y, Li D, He J, Xiao Y (2023) Low-level cadmium exposure induced hormesis in peppermint young plant by constantly activating antioxidant activity based on physiological and transcriptomic analyses. Front Plant Sci 14:1088285. 10.3389/fpls.2023.1088285

Waters ER (2013) The evolution, function, structure, and expression of the plant sHSPs. J Exp Bot 64:391–403. 10.1093/jxb/ers355

Waters ER, Vierling E (2020) Plant small heat shock proteins–evolutionary and functional diversity. New Phytol 227:24–37. 10.1111/nph.16536

Wu Z, Yang L, Jiang L, Zhang Z, Song H, Rong X, Han Y (2019) Low concentration of exogenous ethanol promoted biomass and nutrient accumulation in oilseed rape (Brassica napus L.). Plant Signal Behav 14:1681114. 10.1080/15592324.2019.1681114

Xiao Z, Rogiers SY, Sadras VO, Tyerman SD (2018) Hypoxia in grape berries: the role of seed respiration and lenticels on the berry pedicel and the possible link to cell death. J Exp Bot 69:2071–2083. 10.1093/jxb/ery039

Zhang, JH, Huang, WD, Liu, YP, Pan QH (2005) Effects of Temperature Acclimation Pretreatment on the Ultrastructure of Mesophyll Cells in Young Grape Plants (Vitis vinifera L. cv. Jingxiu) Under Cross-Temperature Stresses. J Integ Plant Biol 47:959−970. 10.1111/j.1744-7909.2005.00109.x

